# Variation and interaction of distinct subgenomes contribute to growth diversity in intergeneric hybrid fish

**DOI:** 10.1101/2024.03.07.584006

**Authors:** Li Ren, Mengxue Luo, Jialin Cui, Xin Gao, Hong Zhang, Ping Wu, Zehong Wei, Yakui Tai, Mengdan Li, Kaikun Luo, Shaojun Liu

## Abstract

Intergeneric hybridization greatly reshapes regulatory interactions among allelic and non-allelic genes. However, their effects on growth diversity remain poorly understood in animals. In this study, we conducted whole-genome sequencing and mRNA-seq analyses in diverse hybrid varieties resulting from the intergeneric hybridization of goldfish (*Carassius auratus* red var.) and common carp (*Cyprinus carpio*). These hybrid individuals were characterized by distinct mitochondrial genomes and copy number variations. Through a weighted gene correlation network analysis, we identified 3693 genes as candidate growth-regulated genes. Among them, the expression of 3672 genes in subgenome R (originating from goldfish) displayed negative correlations with growth rate, whereas 20 genes in subgenome C (originating from common carp) exhibited positive correlations. Notably, we observed intriguing patterns in the expression of *slc2a12* in subgenome C, showing opposite correlations with body weight that changed with water temperatures, suggesting differential interactions between feeding activity and weight gain in response to seasonal changes for hybrid animals. In 40.31% of alleles, we observed dominant *trans*-regulatory effects in the regulatory interaction between distinct alleles from subgenomes R and C. Integrating analyses of allelic-specific expression and DNA methylation data revealed that the influence of DNA methylation on both subgenomes shapes the relative contribution of allelic expression to the growth rate. These findings provide novel insights into the interaction of distinct subgenomes that underlie heterosis in growth traits and contribute to a better understanding of multiple-allele traits in animals.

## Introduction

Hybridization and polyploidization could rapidly shape various genotypes and phenotypes, providing us with abundant materials for studying the contribution of genetic regulation to phenotypes [1]. Interspecific hybridization in some plants, including *Triticum aestivum* × *Secale cereal* [2] and *Brassica nigra* × *B. rapa* [3], is always used to obtain varieties with excellent economic traits. In fish breeding, interspecific hybridization involving different genera or subfamilies has been detected in cyprinid fish [4, 5], salmonid fish [6], and cichlid fish [7]. Among these fishes, *Cyprinus carpio* (common carp) and *Carassius auratus* red var. (goldfish) shared a common whole genome duplication (WGD) event (13.75 million years ago, Mya) as different genera of the subfamily Cyprinidae and then diverged at 10.0 Mya [8]. The specific WGD results in bigger genome sizes and more chromosome numbers (2n: 100) in them than in most carp [9]. Recent studies show that the high genome plasticity and diverse allelic expression shape distinct morphological characters (e.g., body size and color) in some varieties of them, including goldfish and koi carp [10, 11]. Meanwhile, these characteristics contribute to their adaptability in the diverse environment of slow-moving rivers, lakes, and ponds [10-13]. Interesting, a nascent allopolyploid lineage (4nR_2_C_2_, F_3_-F_28_) was successfully established by the hybridization of female goldfish and male common carp and subsequent whole genome duplication [5, 14]. Gene conversion, accompanied by allopolyploidization and multigenerational inheritance, resulted in the emergence of diverse growth phenotypes in the allopolyploid progenies [15, 16].

Body growth, a classic quantitative trait that includes height in humans and weight in domestic animals [17, 18], exhibits significant diversity in fishes. Although genetic variations in individual growth-regulated genes may impact the growth phenotype [18, 19], the rapid genomic variation induced by hybridization and/or polyploidization is considered the most common and swift way of changing this phenotype in nature. Describing a phenomenon known as “heterosis,” if hybrid offspring exhibit faster growth rates and surpass the size of their parents, this trait is utilized to enhance agricultural production [20-23]. Researchers have extensively explored the genetic basis of heterosis, proposing three classic quantitative genetic hypotheses: dominance, overdominance, and epistasis. Moreover, some studies suggest that the emergence of heterosis may be linked to various molecular regulatory mechanisms, including genomic recombination [24], novel epigenetic modifications [25], and alterations in gene expression due to *trans*-regulatory factors from distinct species [26]. Additionally, research has indicated that the differential expression of alleles from different species is influenced by sequence differences in their regulatory regions [27]. In the case of orthologous genes from different genera, the greater sequence differences in their regulatory regions, as compared to intra- and interspecific hybridization, will influence the reshaping of allele-specific expression (ASE) and its impact on the growth phenotype in their hybrid offspring. This aspect promises to be intriguing work.

To explore the impact of copy number variation and mitochondrial regulation on ASE and growth traits, we collected 160 individuals representing six hybrid varieties derived from the intergeneric hybrid lineages of goldfish (2nRR) and common carp (2nCC). The integrated analyses of genomic, DNA methylation, and gene expression data provide a crucial foundation for our research. Our study will expand our understanding of gene interactions and their impact on phenotypes in animals.

## Results

### Determination of genotypes and growth phenotypes among hybrid varieties

Six hybrid varieties, comprising subgenomes R (originating from goldfish) and C (originating from common carp), were collected from individuals aged 8 months and 24 months after hatching, respectively (Figure 1A and Figure S1). To determine the ploidy levels of all hybrid individuals, flow cytometry was employed, using diploid goldfish (2n = 100) as the control group. To identify the source of mitochondrial genomes in reciprocal F_1_ hybrids and allotriploid individuals with the same ploidy level, a fragment of the *cytb* gene in these hybrid individuals was obtained using Sanger sequencing. Subsequently, we compared the obtained sequences with those of goldfish and common carp to determine the origin of their mitochondrial genomes. Moreover, both whole-genome sequencing (WGS) and mRNA-seq data analyses were conducted to validate the genotypes of the six hybrid varieties. Based on the genomic data, the average depth of mapped reads of subgenomes R *vs.* C was approximately 1:1 in 2nCR, 1:2 in 3nRC_2_ and 3nC_2_R, and 2:1 in 3nCR_2_ and 3nR_2_C (Tables S1–S3). Interestingly, we found that fish with identical subgenome ratios showed consistent distributions in the expression values of alleles R *vs.* C, as observed in the mRNA-seq data (Figure 1B). Three clusters of gene expression profiles (cluster 1: 2nRC and 2nCR, cluster 2: 3nR_2_C and 3nCR_2_, cluster 3: 3nC_2_R and 3nRC_2_) remained stable across individuals (10–42 individuals in each variety, Figure 1B). Similarly, analysis of the mRNA-seq data clearly identified mitochondrial types (originating from goldfish or common carp) based on the different number of reads mapped to their two mitochondrial genomes, respectively (Tables S4–S5). The above results can assist us in the initial identification of the genotype among the hybrid varieties and provide insights into their breeding strategies (Figure 1A).

**Figure 1.**
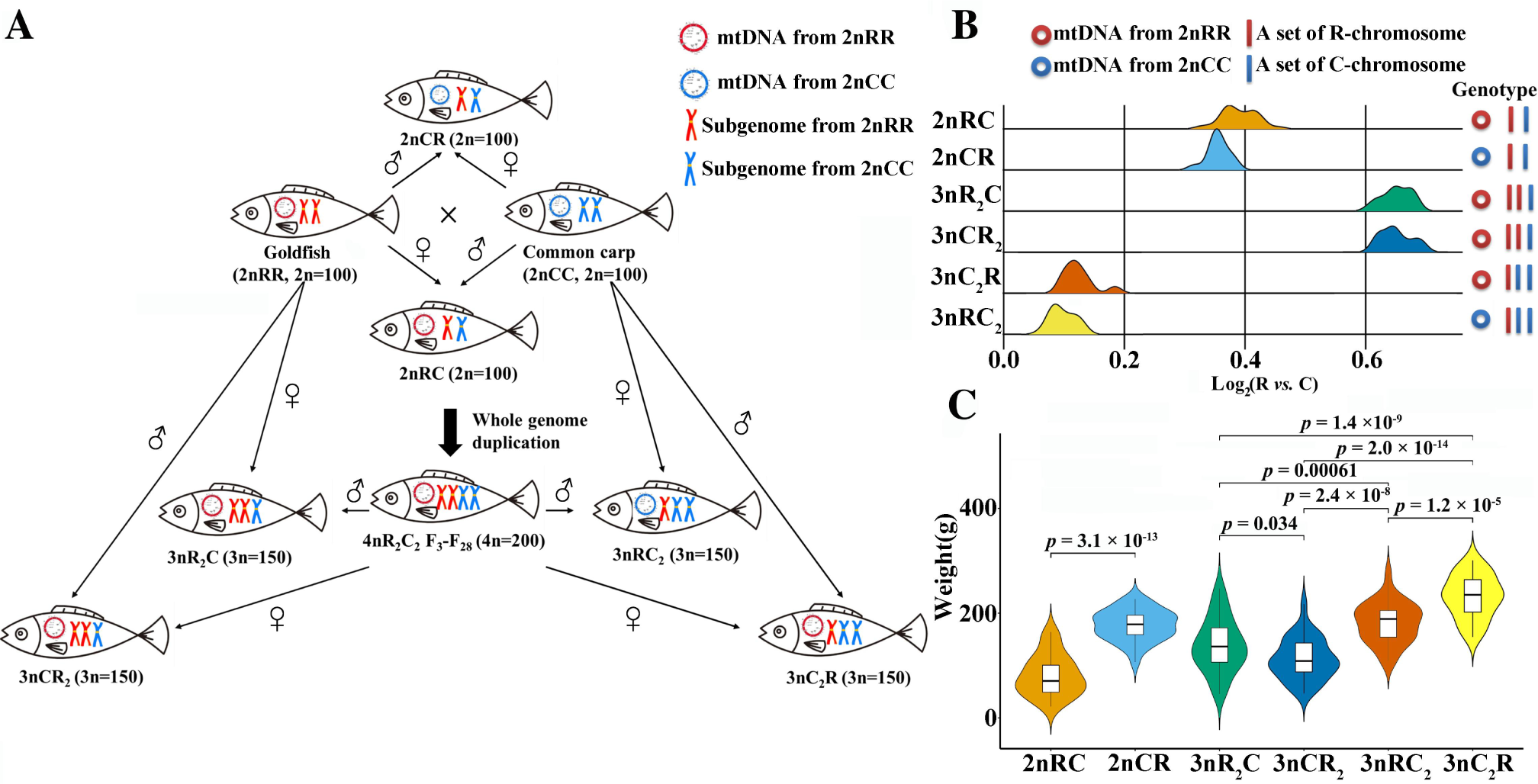
Generated process, genotypes, and growth phenotypes in the six hybrid varieties derived from the intergeneric hybridization of goldfish and common carp. **A.** The controlled genotypes of hybrid varieties obtained from the intergeneric hybridization of goldfish (*C. auratus* red var.) and common carp (*C. carpio*), subsequent polyploidization, and interploid hybridization. **B.** Genotypes of the two interspecific hybrids (2nRC and 2nCR) and the four triploid varieties (3nR_2_C, 3nCR_2_, 3nRC_2_, and 3nC_2_R) predicted based on the mapped reads of transcriptomes. **C.** Body weight in the six hybrid varieties (24 months after hatching).

Analyses of body length (BL), body height (BH), height of back muscle (HBM), and body weight (BW) at 24 months after hatching showed that the growth rate was significantly higher in 2nCR compared to 2nRC (T-test; *P* = 2.08×10^-24^; two-tailed; *t* = 2.02; df = 42, Figure 1C). Meanwhile, the allotriploid 3nC_2_R and 3nRC_2_ (subgenomes R *vs.* C = 1:2) displayed faster growth rates compared to 3nCR_2_ and 3nR_2_C (subgenomes R *vs.* C = 2:1) (T-test; *P* = 4.13×10^-12^; two-tailed; *t* = 1.98; df = 56, Figures 1 and 2). However, no significant difference in growth rate was observed between two fish with the same genotype (3nCR_2_ and 3nR_2_C). Furthermore, a higher growth rate was detected in 3nRC_2_ (mitochondrial genomes originating from common carp) than in 3nC_2_R (mitochondrial genomes originating from goldfish) (T-test; *P* = 0.0001; two-tailed; *t* = 2.01; df = 46, Figure 1C), while there is the same nuclear genome in them (Figure 1A). These findings indicate that more sets of subgenome C and mitochondrial genomes originating from 2nCC contribute to the large body size or rapid growth rate among these hybrid varieties.

**Figure 2.**
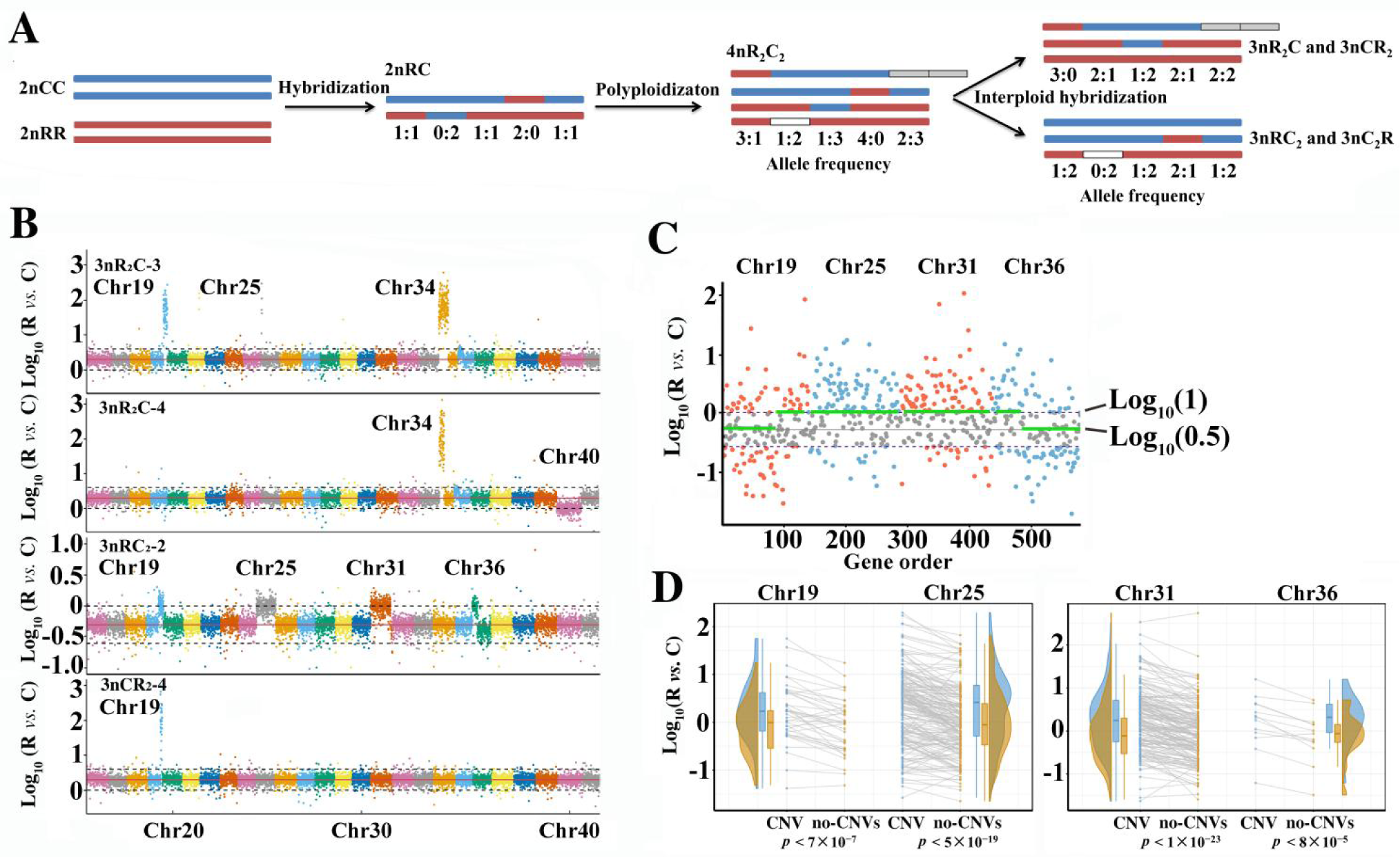
Copy number variations (CNVs) regulating allelic expression changes in the hybrid varieties. **A.** Schematic diagrams of CNVs accompanied by hybridization, polyploidization, and interploid hybridization. White block represents the deletion of allele R or C. Gray box represents the duplication of allele R or C. **B.** Allelic copy number variation resulting from the ratio changes of alleles R *vs.* C. The variation of allelic copy number in the contiguous genes of chromosomes in four triploid individuals. For example, the loss of allele C was observed in the 92 contiguous genes of chromosome 19 (chr19). The 1:1 ratio of allelic copy number was observed in chr40 of 3nR_2_C-4 and in chr19, chr36, chr25, and chr31 of 3nRC_2_-2. In 3nR_2_C and 3nCR_2_, red line represents Log_10_(2), while in 3nRC_2_, it is Log_10_(0.5). Dotted line represents the calibration line, which is used to determine whether CNV occurs. **C.** The changes in the expression ratio of alleles R *vs.* C (chr19, chr25, chr31, and chr36) accompanied by CNVs in 3nRC_2_-2. Green line represents the average values in the no-CNV and CNV regions. **D.** Individuals with no-CNV and CNV had significantly different expression ratios of alleles R *vs*. C (the chr19, chr25, chr31, and chr36 of 3nRC_2_-2) (two-sided T-test analysis).

### High copy number variations in subgenome R

Previous studies demonstrated that gene conversion occurred in the assembled genome of allotetraploid fish [15]. The diverse copy number variations (CNVs) between somatic and germ cells were found in the interspecific F_1_ and allotetraploid populations, revealing various mitotic and meiotic CNVs associated with gene conversion [5, 15]. To investigate the presence of different CNVs in allotriploid progenies, obtained from the backcrossing of allotetraploid to diploid goldfish or common carp, we conducted WGS on seven individuals of the F_1_ hybrid and 22 individuals of triploids (8 months after hatching) (Tables S1–S2). CNV detection for 1545–3017 genes in the 2nRC and triploid varieties revealed a higher CNV ratio in subgenome R (49.25%–83.33%) than in subgenome C (16.67%–50.75%) (T-test; *P* = 4.02×10^-21^; two-tailed; *t* = 2.00; df = 56), indicating a leading role of subgenome R in the genome plasticity of hybrid varieties (Figure 2A and Table S6). Focusing on 46,424 species-specific genes (SSGs) (24,283 in 2nCC and 22,141 in 2nRR) and 18,020 allelic gene pairs (AGPs), the majority of genes with CNVs (71.55%–85.55%) belonged to SSGs and exhibited a higher ratio of genome variation in SSGs than in AGPs (T-test; *P* = 1.80×10^-53^; two-tailed; *t* = 2.00; df = 56) (Tables S6 and S7).

We investigated the distribution of copy number variations (CNVs) in allelic genes. We observed a higher frequency of copy number increase events in allele R (52.65%–74.43%) than in allele C (25.57%–47.35%) (T-test; *P* = 1.04×10-43; two-tailed; *t* = 2.05; df = 28) (Table S7). In some individuals, CNV events occurred on long chromosomal segments involving contiguous genes or entire chromosomes (Figure S2). For example, CNV events occurred on the entire chromosome 40 (chr40) in the 3nR_2_C-4 individual. In the 3nRC_2_-2 individual, CNV events occurred on the parts of chr19 (92 genes) and chr36 (63 genes), as well as the entire chr25 and chr31. These CNV events resulted in a 1:1 ratio of gene copy numbers from the inbred parents in these allotriploid individuals (Figure 2B and Figure S2).

Furthermore, WGS data revealed allele loss caused by CNVs. Among the 22 allotriploid individuals, the number of allele loss events ranged from 2 to 223, while none were observed in the F_1_ hybrids (Table S7). For instance, the loss of allele C in contiguous genes on chr19 was observed in two individuals of 3nR_2_C and one individual of 3nCR_2_ (Figure 2B and Figure S2). We speculated that the shared allele loss observed in different triploid progenies may be derived from the gametes of the same paternal 4nR_2_C_2_ individual through interploid hybridization. Interestingly, the loss of allele C was detected in the 92 contiguous genes of 3nCR_2_-4, which also exhibited the highest growth rate among the 3nCR_2_ population. GO analysis of these 92 genes identified *rfx3* and *gpat4* as being annotated to epithelial cell maturation (GO: 0002071, *p*-value = 0.001). We speculate that the loss of allele C in these two genes may contribute to the high growth rate observed in the allotriploid.

### CNVs altering allele-specific expression

To investigate the impact of CNVs on allelic expression, we conducted integrated genomic and expression analyses using muscle tissue samples from 7 individuals of the F_1_ hybrid and 22 individuals of the allotriploids (Table S2). Specifically, we focused on the loss of allele C in 3nR_2_C and 3nCR_2_, comparing the gene expression values between no-CNV (no allelic loss) and those with CNV (allelic loss) within the corresponding population. In the 3nRC_2_-2 individual, where CNVs led to a 1:1 allelic copy number ratio (Figure 2B), genes within the CNV region also displayed a 1:1 allelic expression ratio (Figure 2C). Conversely, genes outside the CNV regions exhibited a 1:2 allelic expression ratio (Figure 2C). Then, differential expression analysis was performed between CNV and no-CNV individuals in the 3nRC_2_ population, and significant differences were detected in chr19, chr25, chr31, and chr36 (Figure 2D). These findings suggest that copy number changes in the hybrids could affect gene expression, highlighting the effects of CNVs on allelic expression.

### Mitochondrial regulation shaping allele-specific expression and species-specific expression

Distinct mitochondrial genomes and the same nuclear genome were in two groups (diploid group 1: 2nRC and 2nCR, triploid group 2: 3nRC_2_ and 3nC_2_R). This provided valuable insights into mitochondrial genetics and its role in regulating growth diversity through gene expression (Tables S1–S2 and S8). Comparative analyses revealed that 6516 genes (7.9%) of the total 82,464 genes (alleles R and C considered independent genes) differed in expression between the reciprocal F_1_ hybrids (13 individuals in 2nRC and 2nCR), while only 126 genes (0.15%) showed differential expression between 3nRC_2_ and 3nC_2_R (10 individuals). The lower number of differentially expressed genes (DEGs) in triploids compared to diploids may be caused by the high frequency of CNVs disrupting allele-specific expression (ASE) regulated by mitochondrial genetics (Figure 3A).

**Figure 3.**
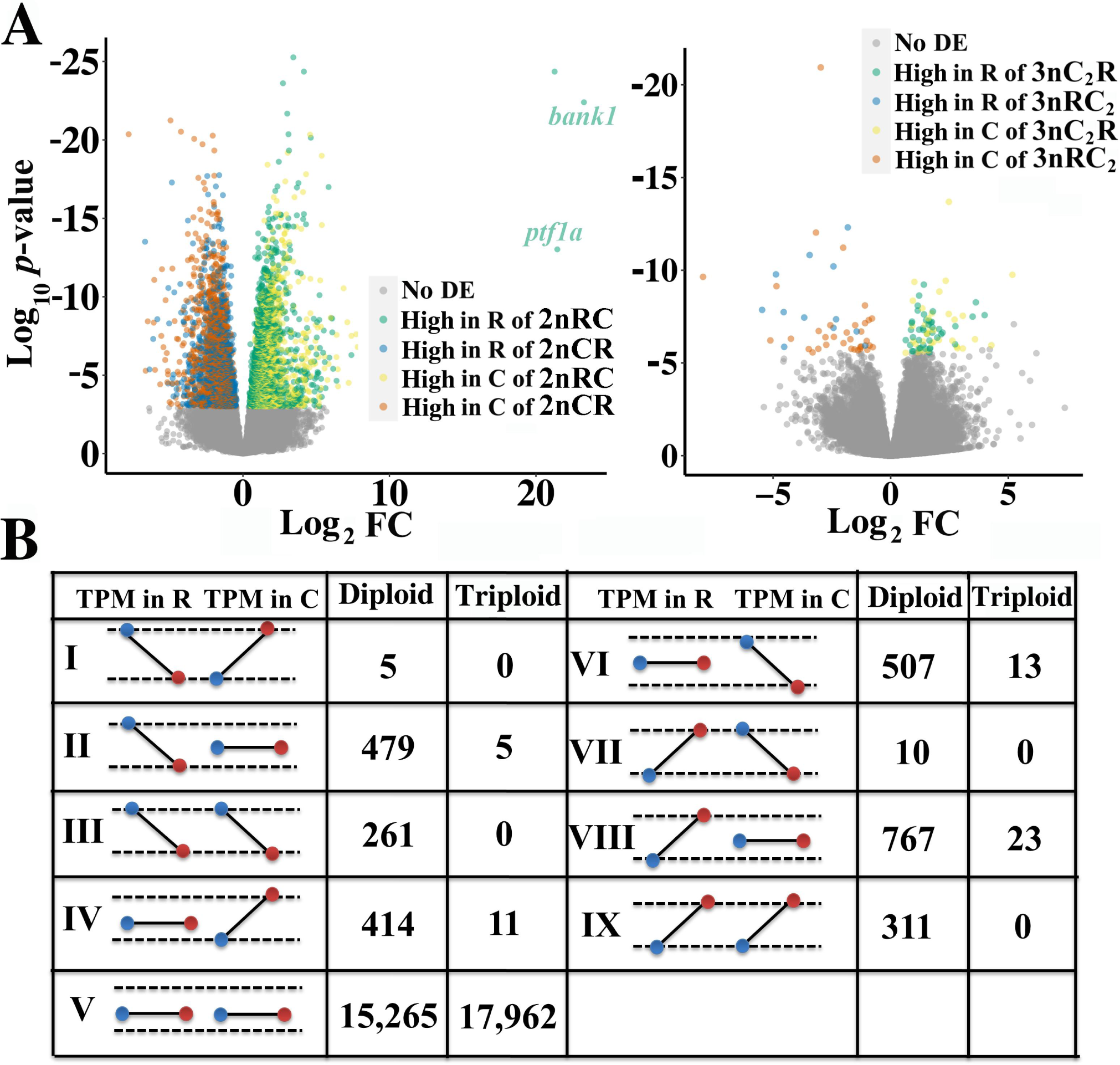
Mitochondrial genetics regulating allele-specific expression in reciprocal hybrids. **A.** Differential expressed analysis was performed in the diploid (2nRC and 2nCR) and triploid (3nC_2_R and 3nRC_2_) groups. “no DE” represents no differential expression; “high in R of 2nRC” represents the genes relating to higher expression in allele R of 2nRC than ones in 2nCR (1881 genes, green); “high in R of 2nCR” (1393 genes, blue); “high in C of 2nRC” (1545 genes, yellow); “high in R of 2nCR” (1697 genes, orange); the silencing of allele R was detected in the two genes (*bank1* and *ptf1a*) of 2nCR, while gene expression was observed in alleles of 2nRC. **B.** Schematic diagrams of nine allelic expression patterns regulated by mitochondrial genetics. Blue represents the expression values in 2nCR and 3nRC_2_ (both mtDNA originating from both 2nCC), while red represents the expression values in 2nRC and 3nC_2_R (both mtDNA originating from both 2nRR). “TPM in R” represents the expression values in allele R.

In the diploid group, the analysis of differential gene expression revealed that in subgenome R, AGPs (1834) had more DEGs compared to SSGs (1440), whereas in subgenome C, SSGs (1734) had more DEGs than AGPs (1508) (Pearson’s chi-squared test; *P* = 2.04×10^-14^; χ^2^ = 58.497; df = 1) (Table S9). This suggests that orthologous genes from goldfish and species-specific genes from common carp are more susceptible to regulation by maternal effects, resulting in differential expression between reciprocal F_1_ hybrids. Furthermore, we detected that highly expressed genes in 2nRC were predominantly in allele R (1881) rather than in allele C (1545), while the opposite trend was observed in 2nCR, where highly expressed genes were more in allele C (1697) than in allele R (1393) (Figure 3A). This finding suggests that mitochondria in hybrids may preferentially upregulate the expression of nuclear genes from the same species. Interestingly, we noticed that allele R in *ptf1a* was silenced in 2nCR, whereas its expression was detected in 2nRC, indicating that the mitochondrial genes originating from 2nCC inhibited the expression of allele R (Figure 3A). The significant diversity in allelic gene expression between 2nRC and 2nCR, resulting from the silencing of allele R in *ptf1a*, may be linked to their distinct growth rates [28].

To further investigate the magnitude of independence and interaction between alleles R and C, we established nine expression patterns for alleles R and C (Figure 3B). Out of these, 2754 genes (15.28%) in the diploid group (2nRC and 2nCR) and 52 genes (0.28%) in the triploid group exhibited expression changes regulated by distinct mitochondrial genetics. In the diploid group, we found that the expression of either allele R or C was altered in specific patterns (II, IV, VI, and VIII) across 2167 genes (12.02%). These patterns indicate the existence of independent regulatory networks connecting mitochondrial genes to the expression of alleles R or C (Figure 3B and Figure S3). Additionally, a shared regulatory network in mitochondrial genetics regulating both alleles R and C (patterns III and IX) was observed in 572 genes (3.17%) (Figure 3B and Figure S3). Interestingly, we observed an opposite trend between the expression of alleles R and C in 15 genes (0.08%, patterns I and VII), which was likely related to an antagonistic relationship between the regulatory networks of the distinct alleles. These results showed the diversified regulatory networks in mitochondrial genetics that regulate allelic gene expression.

### Gene coexpression analyses revealing genetic networks underlying growth rate diversity

The above results demonstrated that the diversified ASE and species-specific expression were in the six hybrid varieties. We then used a weighted gene correlation network analysis to detect gene coexpression networks and genetic modules in the 131 individuals (24 months after hatching). Twelve modules were identified based on the correlation of the expression profiles of the 65,495 expressed genes (31,974 genes in subgenome R, 33,521 genes in subgenome C) (Figure S4A). Among these modules, Module Eigengene 1 (ME01) exhibited the highest correlation with growth-related phenotypic values, including body weight and body length, while a significant correlation was detected between gene significance and module membership (*r* = 0.81, *p* = 1 × 10^-20^, Figure S4B). After gene filtering, we identified 3693 genes within ME01 as candidate growth-regulated genes. Within this module, the coexpression network reveals that the 3672 genes in subgenome R and the 21 genes in subgenome C showed significant correlations between gene expression and body weight (Figure 4A). These results indicate that CNVs in subgenome R, along with changes in gene expression within subgenome R, are primarily responsible for regulating growth diversity in these hybrid varieties.

**Figure 4.**
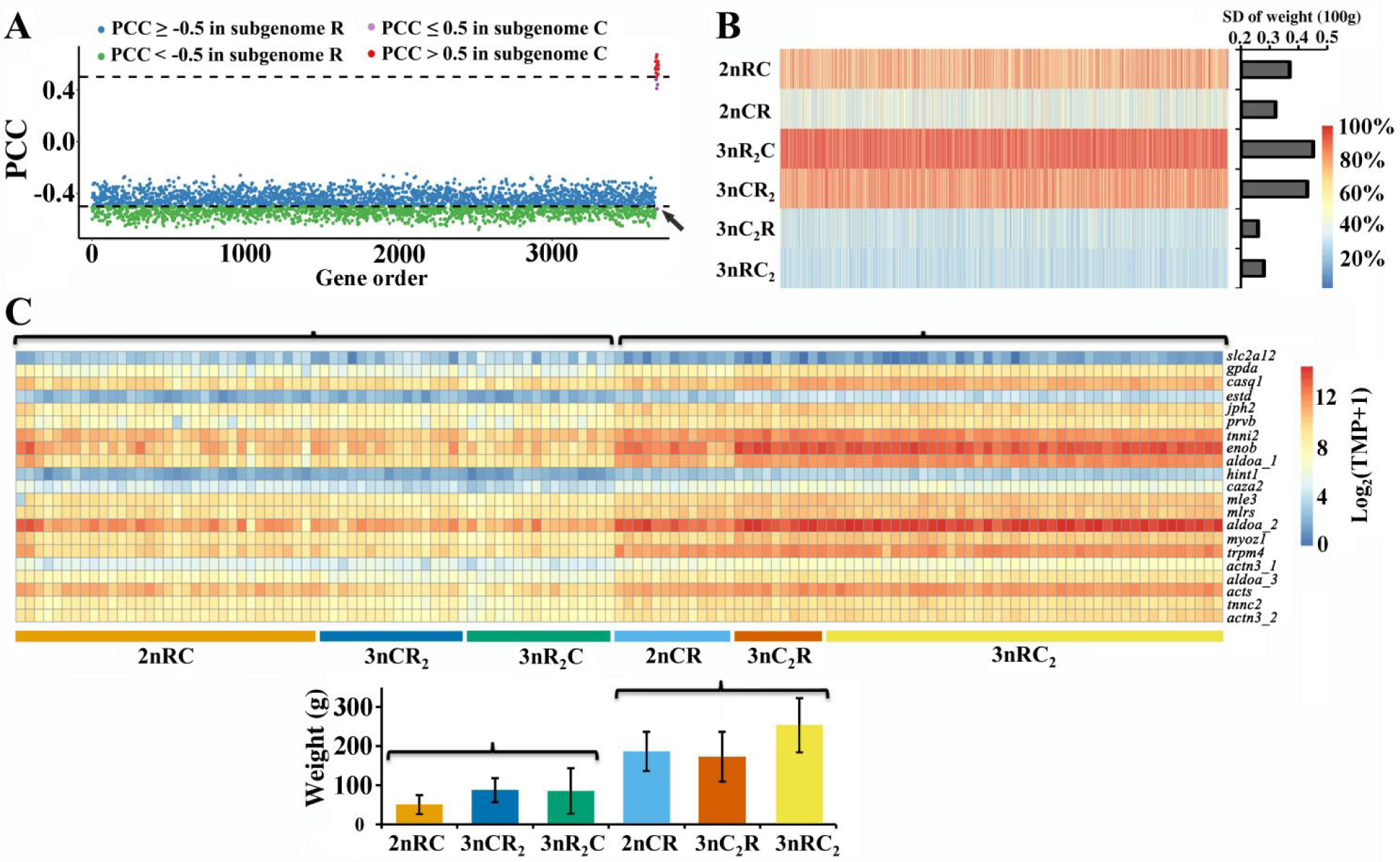
Expression of 3693 growth-regulated genes altering body weight in the hybrid varieties. **A.** Pearson correlation coefficient (PCC) between the values of gene expression and body weight in the 3693 predicted growth-regulated genes. Among these genes, 3672 genes originated from subgenome R. Among them, the PCC values of 2032 genes are represented as blue dots and are greater than or equal to -0.5. Meanwhile, the PCC values of 1640 genes are represented as green dots and are less than -0.5. A gene (Solute Carrier Family 2 Member 12, abbreviated as *slc2a12*, black arrow) in subgenome C exhibited a negative correlation between gene expression and body weight (24 months after hatching). The other 20 genes in subgenome C showed positive correlations. **B.** Diversities in gene expression and body weight across the individuals of each variety. The standard deviation (SD) of GPM values in the 3693 growth-regulated gene was performed to assess the expression diversity across individuals. The SD of body weight values was calculated to assess the growth diversity across individuals. **C.** A heatmap exhibiting the expression of the 21 growth-regulated genes in subgenome C across individuals. Two groups were classified into the six hybrid varieties based on the clustering of body weight and gene expression.

To further identify the relationship between the expression of the 3693 growth-regulated genes and their body weight, we analyzed gene expression diversity and body weight diversity across individuals. The highest diversities of gene expression and body weight were observed in 3nR_2_C, while the lowest diversities were observed in 3nC_2_R (Figure 4B). Importantly, both 3nR_2_C and 3nCR_2_ (with two sets of subgenome R) exhibited higher diversities in gene expression and body weight compared to 3nRC_2_ and 3nC_2_R (with one set of subgenome C). These findings, coupled with the enrichment of CNVs in subgenome R, indicate that the high CNV ratio in subgenome R contributes to the increased diversity of allele R expression, which subsequently leads to the diversification of growth phenotypes in the hybrid population (Figure 2 and Table S6). In the diploid group, higher gene expression diversity was detected in 2nRC than in 2nCR (Figure 4B), suggesting that the maternal effects related to goldfish-originated regulation may be more beneficial to the CNV of allele R and result in gene expression diversity across individuals.

Among the 3693 growth-regulated genes, we found that the expression of 3672 genes in subgenome R and one gene (*slc2a12*) in subgenome C exhibited negative correlations with body weight, while the other 20 genes in subgenome C showed positive correlations (Figure 4A). Additionally, the expression of these 20 genes was found to be higher in the 2nCR, 3nRC_2_, and 3nC_2_R populations compared to the 2nRC, 3nR_2_C, and 3nCR_2_ populations (Figure 4C). We compared gene expression data from 24-month-old individuals (n = 132) with data from 8-month-old individuals (n = 29). Genes exhibiting strong positive correlations (Pearson correlation coefficient, PCC > 0.5) between body weight and gene expression in the larger dataset also showed strong positive correlations in the smaller dataset, with the exception of the *prvb* gene (PCC = 0.37, *p*-value = 0.05) (Figure S5). Interestingly, the expression of *slc2a12* in subgenome C exhibited a positive correlation with body weight at low water temperature (about 8 ℃, 8 months after hatching) (PCC = 0.67, *p*-value = 0.003) (Figure S5), while a negative correlation was detected at high water temperature (about 20 ℃, 24 months after hatching) (Figure 4, A and C). The opposite correlations were likely related to the different strategies for rapid growth rate of these hybrid individuals in alternation with the seasons, in which the up-regulated expression of *slc2a12* in subgenome C could decrease the amount of exercise and energy consumption at low temperatures (winter), while the down-regulated expression could increase the amount of exercise for obtaining food at high temperatures (spring) [29].

### Variations in gene regulatory networks and their effects on growth rate

The decreased expression of genes in subgenome R and the increased expression of genes in subgenome C both contributed to the rapid growth rate, prompting the question of how variations in gene regulatory networks regulated allelic expression and growth diversity. Among the 3693 growth-regulated genes, 2094 were orthologous genes, while 1587 were specie-specific genes (SSGs) in subgenome R and 12 were specie-specific genes (SSGs) in subgenome C (Figure S6). The PCC values of the 3693 growth-regulated genes were higher in the 2094 AGPs than in non-AGPs (ANOVA *F*-test = 0.85, df = 2081, *p* = 0.000123) (Figure 5A). This finding suggests that shared *trans*-regulatory factors between alleles R and C homogenize the expression levels of the two orthologous genes originating from goldfish and common carp, leading to increased synchrony in the allelic expression in hybrids. In the analysis of PCC between 2094 AGPs, 832 (39.72%) displayed a positive correlation (PCC > 0.3), with 304 (14.52%) exhibiting a strong positive correlation (PCC > 0.5). In contrast, only 12 AGPs (0.57%) displayed a negative correlation (PCC < -0.3), and no AGPs exhibited a strongly negative correlation (PCC < -0.5). These results reflect a synchrony of allelic expression in 40.31% of genes and independence in 59.69% of genes. Furthermore, we analyzed nine AGPs where the expression of allele C exhibited a strong positive correlation (PCC > 0.5) with body weight (Figure 5B). Among these nine AGPs, six also showed a strong positive correlation (PCC > 0.5) between the expression values of alleles R and C (Figure 5B). Interestingly, higher PCC values of two alleles and lower body weights were observed in 2nRC than in 2nCR, and the same phenomenon was detected in 3nR_2_C and 3nCR_2_ than in 3nRC_2_ and 3nC_2_R (Figure 5C). Among hybrids with the same ploidy level, more diversified gene regulatory networks between alleles of growth-regulating genes may be associated with faster growth. This result sheds light on why heterosis often appears in hybrid F_1_ generations, and with successive generations of hybrid offspring, the heterosis diminishes or decreases [20-24]. This phenomenon might be attributed to the presence of complete *trans*-regulatory factors from different species only in hybrid F_1_, leading to the maximization of differences in the gene regulatory network between allelic genes.

**Figure 5.**
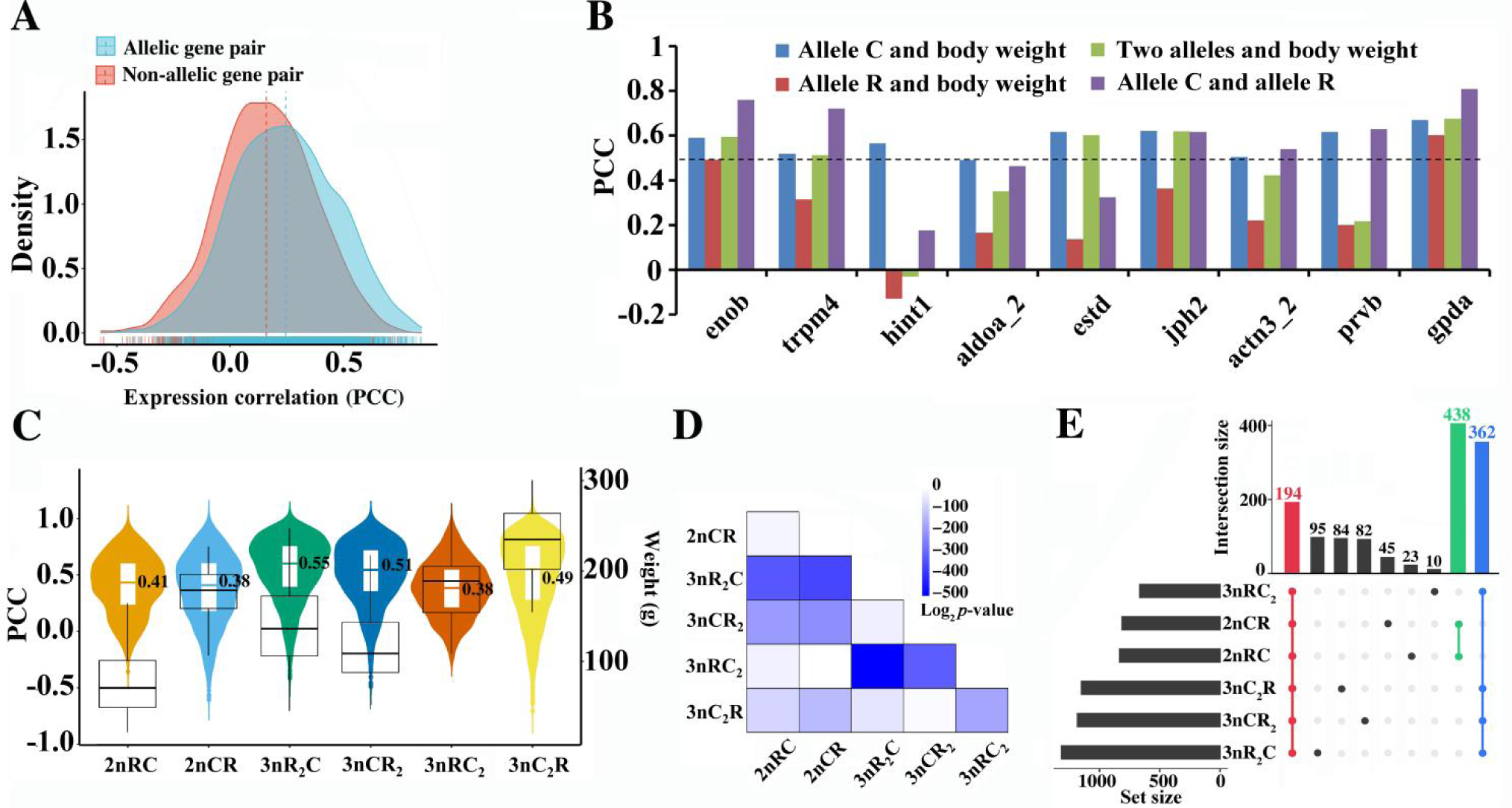
Correlational analyses of expression of alleles R and C, allele-specific expression, and body weight. **A.** Expression correlational analyses of the 3693 growth-regulated genes between the expression of AGPs and non-allelic gene pairs based on Pearson correlation coefficient (PCC) values. **B.** Correlational analyses of the nine AGPs, which belong to the 21 growth-regulated genes in subgenome C. Blue represents the Pearson correlation coefficient between the expression of allele C and body weight. **C.** The distributions of PCC (violin plot) and body weight (box plot) values in the six hybrids. The median value of PCC is indicated in Figure. **D.** A two-sided *t*-test of PCC values was performed to detect the effects of gene regulatory networks between two hybrid varieties. **E.** The distribution of genes with |PCC| > 0.5. For example, 194 genes were shared among the six hybrid varieties (red).

The PCC values between alleles could also allow us to assess the variation of gene regulatory networks among the six hybrid varieties. The highest difference was observed between 3nR_2_C and 3nRC_2_ (T-test; *P* = 6.9 × 10^-152^; two-tailed; *t* = 1.96; df = 2084) (Figure 5D), which had distinct mitochondrial genomes and different subgenome ratios of R *vs.* C (Figure 1A). We further investigate the differences in gene regulatory networks influenced by mitochondrial genetics in the two reciprocal F_1_ hybrids (T-test; *P* = 2.66 × 10^-6^; two-tailed; *t* = 1.96; df = 2084) (Figure 5D). Reversed PCC values were observed in 310 growth-regulated AGPs (Table S10). Further analysis of the strong positive correlation (PCC > 0.5) between the expression of two alleles revealed that only 194 AGPs (10.35% of 1874 AGPs) were shared among the six hybrid varieties, while 439 common AGPs were observed between the two F_1_ hybrids and 362 common AGPs were identified among the four triploid varieties (Figure 5E and Figure S7). Further, 2nCR and 3nRC_2_ exhibited similar PCC in allelic gene expression (T-test*: P* = 0.80; two-tailed; *t* = 1.96; df = 2084), raising the question of why these hybrids with different ploidy levels exhibited similar gene regulatory networks. Recent studies revealed that DNA methylation inhibits the transcription of genes in the additional subgenome C of 3nRC_2_ (Figure 6A) [30], which may account for the similar gene regulatory networks between 2nCR and 3nRC_2_. The above results revealed a high divergence of gene regulatory networks between alleles in these hybrid varieties.

**Figure 6.**
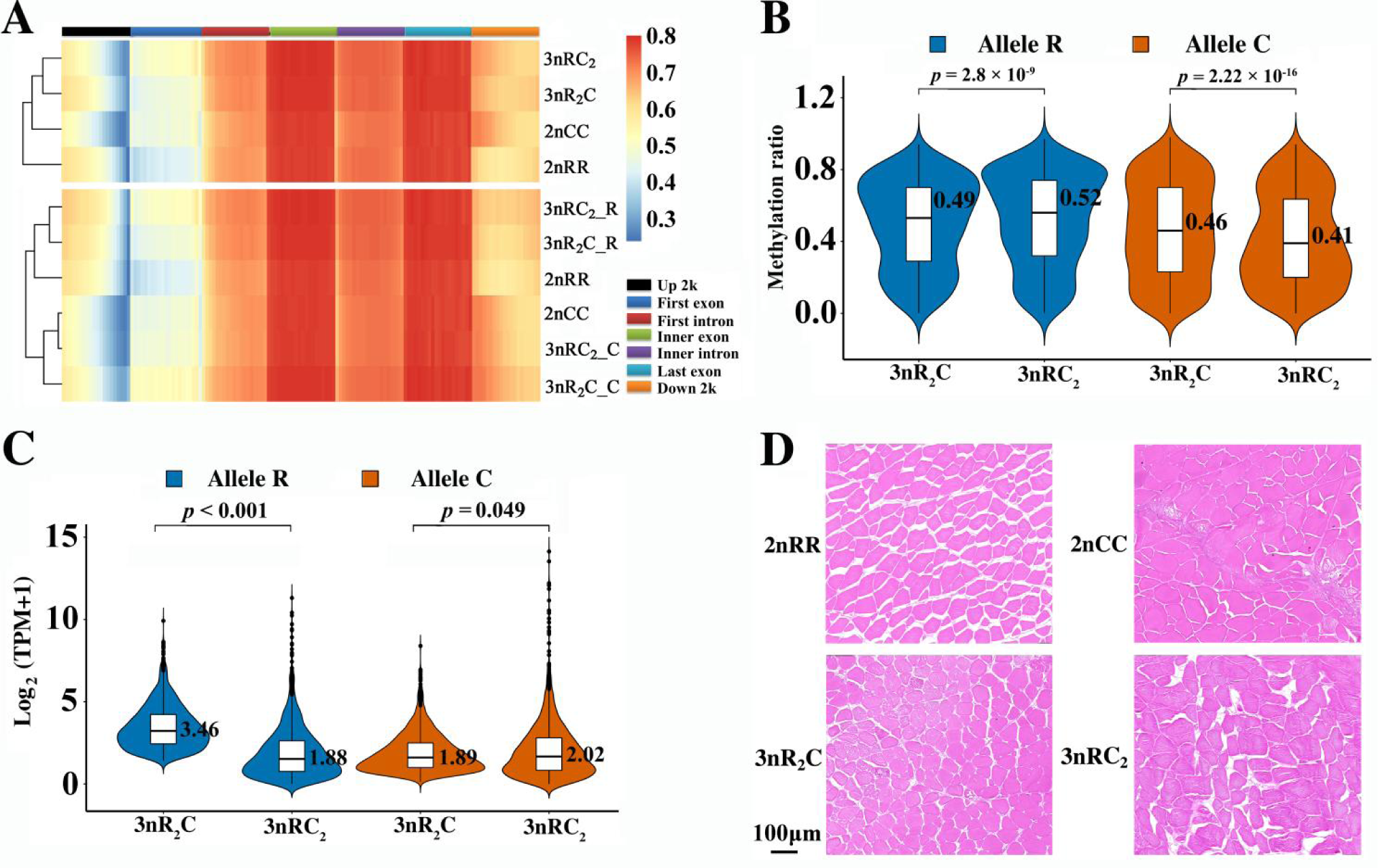
Analyses on DNA methylation, allele-specific expression, and histological observation between hybrids with different body weights. **A.** DNA methylation levels of different gene elements in the two triploids (3nR_2_C and 3nC_2_R) and their inbred diploid parents (2nRR and 2nCC). Each region was divided into twenty bins based on its total length. The methylation ratio of subgenome R was higher in two triploids than in 2nRR, while the methylation ratio of subgenome C was higher in 3nR_2_C than in 3nC_2_R and 2nCC. “3nR_2_C” represents the combined methylation ratios of subgenomes R and C in 3nR_2_C. “3nR_2_C-R” represents the methylation ratios in the subgenome R of 3nR_2_C**. B.** DNA methylation level of the 2094 growth-regulated genes. **C.** The expression levels of the 2094 growth-regulated genes. **D.** Cross-section of skeletal muscle (H&E staining) showing the myofibers of the two triploids and their inbred parents (2nRR and 2nCC) (n = 3 biologically independent samples). Scale bar = 100 μm (10X).

### Allele-specific DNA methylation regulating growth rate diversity

Through analyses of DNA methylation in the 3693 growth-regulated genes, we try to investigate whether and how DNA methylation affects growth rates through regulating ASE. So we analyzed whole-genome bisulfite sequencing data (the muscles of 2nRR, 2nCC, 3nR_2_C, and 3nRC_2_) and growth phenotype data (a significant difference in body weight between 3nR_2_C and 3nRC_2_) to investigate the contribution of DNA methylation to subgenomes R and C [30]. After mapping clean reads to the combined genome of 2nRR and 2nCC, the uniquely mapped reads were used to assess the methylation levels in CpG islands. Comparative analysis revealed that the DNA methylation level in each subgenome was higher in the two triploids (3nR_2_C and 3nRC_2_) than in their inbred parents (2nRR and 2nCC) (Figure 6A), indicating that high DNA methylation suppressed gene expression in triploids.

We focused on 2094 genes that are involved in growth regulation, and we compared the DNA methylation levels of the alleles R and C in 3nR_2_C and 3nRC_2_. We found that allele R showed lower DNA methylation in 3nR_2_C than in 3nRC_2_, while its expression was higher in 3nR_2_C (Figure 6B–C). On the other hand, allele C showed higher DNA methylation in 3nR_2_C and lower expression compared to 3nRC_2_ (Figure 6B–C). Comparing 3nR_2_C and 3nRC_2_, we identified 85 differentially methylated genes (DMGs) (4.06%) from AGPs and 57 DMGs (3.56%) from SSGs (Table S11). The majority of DMGs (AGP: 73 of 85, SSG: 46 of 57) had reduced DNA methylation in 3nR_2_C (Table S11), indicating that the lower DNA methylation in subgenome R contributed to the higher expression levels of allele R in 3nR_2_C compared to 3nRC_2_. Indeed, histological examination of skeletal muscle tissues using hematoxylin and eosins (H&E) staining uncovered that the cross-sectional area of myofibers was bigger in 2nCC than in 2nRR, which was likely related to the difference in skeletal muscle development between the two species (T-test; *P* = 3.18×10^-16^; two-tailed; *t* = 2.00; df = 56) (Figure 6D). We detected a larger cross-sectional area of myofibers in 3nC_2_R than in 3nR_2_C, suggesting that the high expression of allele C benefited the growth of myofibers (T-test; *P* = 1.40×10^-5^; two-tailed; *t* = 2.00; df = 56) (Figure 6D). In conclusion, our findings indicate that the regulation of DNA methylation may contribute to the differential activity of ASE and growth rates between 3nC_2_R and 3nR_2_C, possibly explaining the variations in their growth rates.

## Discussion

Using different hybrid strategies that produce the controlled genotypes of hybrids, we characterized the mitochondrial and nuclear genetic variants associated with variations in gene expression and growth diversity. We detected the CNVs of subgenomes R and C across hybrid individuals, including allele loss in triploids, and found their effects on allele-specific and species-specific expressions. Applying a weighted gene correlation network analysis, we identified 3693 genes as candidate growth-regulated genes, in which the expression of 2094 allelic gene pairs and 1599 species-specific genes exhibited a significant correlation with growth diversity. Using correlation analyses between the expression of distinct alleles R and C, we detected the different degrees of independence and interactions in the allelic regulatory networks, which reflected variations of gene regulatory networks among different hybrid varieties. Importantly, we found that the diversified gene regulatory networks of distinct alleles R and C in growth-regulated genes may contribute to the rapid growth rate. This result suggests that maintaining and increasing differentiation in allelic expression will result in the emergence of heterosis and be beneficial for aquaculture and animal breeding. In addition, we revealed that DNA methylation shaped allelic expression variations in subgenomes R and C.

Following their divergence 10 Mya, high genome plasticity in goldfish and common carp facilitated diverse growth phenotypes during domestication [8, 31, 32]. Now, we detected diverse CNVs in the nascent allotetraploid population (4nR_2_C_2_, F_3_–F_28_) [15, 33], which were derived from the hybridization of goldfish and common carp [5, 14], and provided abundant genetic diversities in their allotriploid progenies through different interploid crossings with their inbred parents [16]. Meanwhile, the random emergence of repair DNA damage during mitosis also resulted in CNVs, including allele loss in triploid individuals. Joint analyses of the CNV region, growth rate, and functional annotation data could provide an effective way to identify causal genes associated with the growth variation in inter- and intra-hybrid populations. For example, we showed that the loss of allele C in *rfx3* and *gpat4* might have contributed to the high growth rate. The majority of CNVs in the hybrid varieties were distributed in subgenome R, where dynamic transposition of transposable elements could result in CNVs and allelic expression variation and increase growth diversity [15].

Diverse subgenome ratios, mitochondrial genetics, and CNVs provide abundant genetic materials for investigating the relative contribution of gene regulatory networks to the variations in allele- and species-specific expressions. When exposed to the common *trans*-acting regulatory factors, the distinct *cis*-regulatory elements in allelic genes may cause a target gene to interact or bind differentially with the transcriptional factors, thus resulting in differential expression between alleles [34, 35]. We showed that the dominance of *trans*-acting regulatory effects decreased the expression diversity of alleles in most genes, while *cis*-regulatory regulatory effects increased it in a few genes [36, 37]. Different *trans*-acting influences involving mitochondrial regulation in the reciprocal F_1_ hybrids reflect differential expression of species-specific genes primarily in subgenome C, which play key roles in growth diversity. Additionally, the distinct subgenome ratios in allotriploids also shape allelic CNVs and alter the dosage of the *trans*-acting factors between alleles, although the diverse effects of DNA methylation occur in different subgenomes [26, 38]. Our findings indicate that dynamic changes in variations of gene regulatory networks increase the magnitude of independence and interactions in allelic regulatory networks, resulting in a great increase in allelic expression variation and growth diversity in these intergeneric hybrid varieties.

A previous study indicated that the expression dominance in allele R was beneficial to high body height in allotriploids, while the expression dominance in allele C was beneficial to high body length [30]. We showed that the decreased expression of genes in subgenome R and the increased expression of genes in subgenome C contributed to the rapid growth rate. These findings suggest that the appropriate combination of allele-specific expression may contribute to the heterosis in quantitative traits. Interestingly, *slc2a12* belongs to a family of transporters that catalyze the uptake of sugars through facilitated diffusion [39]. The increased expression in winter could decrease the amount of exercise and energy consumption in low temperatures, while the decreased expression in spring could increase the amount of exercise needed to obtain food [29]. These different strategies in alternation with the seasons will increase the growth rates of these hybrid individuals. Our result provided new viewpoints: the great diversity in *cis*-regulatory sequences between distinct alleles could decrease the synergy of allelic-specific expression and increase the magnitude of heterosis in the growth rate. In summary, our findings shed light on how the regulatory network underlying distinct subgenomes regulates growth plasticity.

## Conclusions

The allotriploid obtained through interploidy hybridization between allotetraploid and diploid has the advantages of ecological friendliness (sterility) and a fast growth rate, which have made contributions to fish breeding for more than 20 years in China. It has been a huge challenge for us to figure out how to improve the allotriploid fish even more. Our results revealed that variations in CNVs and gene regulatory networks between alleles could shape the growth diversity in the hybrid populations. Consequently, selecting individuals with low-copy growth-regulated genes from allele R (originating from goldfish) and high-copy growth-regulating genes from allele C (originating from common carp) will contribute to the breeding of allotriploid populations exhibiting a rapid growth phenotype. These studies will help us develop a novel hybrid breeding strategy through the genotype selection of parental allotetraploids.

## Materials and methods

### Collection and determination of samples

The fish used in this study included an F_1_ diploid hybrid (2nRC) obtained from hybridization between *C. auratus* red var. (goldfish, 2nRR, ♀) and *C. carpio* (common carp, 2nCC, ♂), an F_1_ diploid hybrid (2nCR) obtained from hybridization between 2nCC (♀) and 2nRR (♂), an allotriploid (3nR_2_C) obtained from interploid crossing of 2nRR (♀) with a allotetraploid of 2nRR × 2nCC (4nR_2_C_2_, ♂), a allotriploid (3nRC_2_) obtained from interploid crossing of 2nCC (♀) with 4nR_2_C_2_ (♂), a allotriploid (3nCR_2_) obtained from interploid crossing of an allotetraploid of 4nR_2_C_2_ (♀) with 2nRR (♂), a allotriploid (3nC_2_R) obtained from interploid crossing of 4nR_2_C_2_ (♀) with 2nCC (♂). These hybrid varieties, including two reciprocal F_1_ hybrids (2nRC and 2nCR) and four triploids (3nR_2_C, 3nRC_2_, 3nCR_2_, and 3nC_2_R), were fed in separate pools under identical environmental conditions. These conditions included suitable water temperatures, oxygen levels, food supply, breeding density, etc. These pools were located in the drainage area of Dongting Lake, Hunan, China (29° 11’ 51” N, 112° 35’ 50” E). Some growth traits, including body length (BL), body height (BH), height of back muscle (HBM), and body weight (BW), were detected in multiple growth stages. Twenty-nine healthy individuals (eight months after hatching, 7 in 2nRC, 4 in 3nR_2_C, 7 in 3nRC_2_, 8 in 3nCR_2_, and 3 in 3nC_2_R) and 131 healthy individuals (24 months after hatching) were collected for this study, respectively. These hybrid varieties were deeply anesthetized with 300 mg/L Tricaine Methanesulfonate (Sigma-Aldrich, St. Louis, MO, USA) for 10 min (25°C) in a separation tank. After confirming the death, all samples were collected for dissection. The DNA content of erythrocytes from 2nRR, 2nCC, and the hybrids was measured using flow cytometry (Cell Counter Analyzer, Partec, Germany) for identifying chromosome number [40].

### DNA isolation and genomic sequencing

High-quality genomic DNA of 2nRR, 2nCC, and the hybrids was isolated from the muscle tissue using DNeasy Blood & Tissue Kits (Qiagen). The quality of DNA was checked by a NanoDrop^®^ ND-1000 spectrophotometer with 260/280 and 260/230 ratios. The type of mitochondrial genome in the hybrids was identified based on a fragment of cytochrome b (*cytb*). Then, the high-quality DNA was used to construct a paired-end library (150 bp × 2) and sequenced by Illumina X Ten platform according to standard protocol. The detail was as follows: a mixture containing equal amounts of 2nRR and 2nCC DNA, the muscle of 29 hybrid individuals. After obtaining the raw data, the sequencing adaptors were removed. Fastp (v. 0.21.0) was used to remove duplicate read pairs and low-quality reads based on the default parameters [41].

### Detection of copy number variation

High-quality reads in the hybrid were mapped to the combined nuclear and mitochondrial genomes of goldfish [8] (nDNA: PRJCA001234 of NGDC database, mtDNA: AY714387.1) and common carp [32] (nDNA of Yellow River carp: PRJNA510861 of NCBI database, mtDNA: AP009047.1) using BWA with the default parameters. Coordinate-sorted BAM output files of WGS were obtained to calculate the number of mapped reads in the coding region of each gene using htseq-count (v. 0.12.4) with thresholds of “-m union --nonunique=none”. The Gene Per Million (GPM) is a value to measure how many reads are mapped to each gene in genomic data. It helps us understand the relative abundance of gene copies in a sample by considering the length of the gene and the total number of reads. The formula for calculating GPM is as follows:

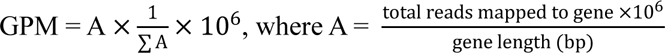

For *in silico* F_1_, we sequenced an equal mixture of 2nRR and 2nCC DNA using the same sequencing platform as other genomic data. By comparing the hybrid varieties with the *in silico* F_1_, we were able to detect copy number variations (CNVs) in all the hybrid varieties. To identify CNVs, we set thresholds based on the genotype. We used the logarithm base 2 of the ratio (GPM_hybrid_/GPM_mixed_) and compared it to 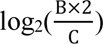 and 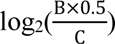. If the logarithm base 2 of (GPM_hybrid_/GPM_mixed_) was greater than 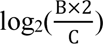 or less than 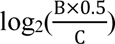 it was considered a CNV. In this formula, “ B “ represents the allelic ratio (R or C) in hybrids, and “C” represents the allelic ratio (R or C) in *in silico* F_1_. for 2nRC, “B” is 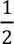; for allele R of 3nRC_2_ and 3nC_2_R, and allele C of 3nR_2_C and 3nCR_2_, “B” is 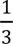; for allele C of 3nRC_2_ and 3nC_2_R, and allele R of 3nR_2_C and 3nCR_2_, “B” is 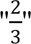.

Allelic gene pairs (AGPs) between the subgenomes R (originating from 2nRR) and C (originating from 2nCC) in the hybrid varieties, including the F_1_ hybrids (2nRC and 2nCR) and four triploids (3nR_2_C, 3nRC_2_, 3nCR_2_, and 3nC_2_R), were obtained using the all-against-all reciprocal BLASTP (v. 2.8.1) with an e-value of 1e ^−6^ based on protein sequences. Then, transcripts that lacked gene annotation and were shorter than 300 bp were discarded from AGPs. GPM values in each AGP could be used to assess allelic CNVs between the subgenomes R and C in the hybrid varieties. The values of log_10_((R_GPM in hybrid_/C_GPM in hybrid_)-(R_GPM in mixed_/C_GPM in mixed_)) could be used to assess the allelic CNVs. The detail thresholds for them were set up based on the genotype as follows: 1) log_10_(2) and log_10_(0.5) in F_1_ hybrid (2nRC and 2nCR); 2) log_10_(4) and log_10_(1) in 3nR_2_C and 3nCR_2_; 3) log_10_(1) and log_10_(0.25) in 3nC_2_R and 3nRC_2_.

### RNA isolation and mRNA-seq

To obtain gene expression profiling of the two reciprocal F_1_ hybrids and allotriploids, total RNA of the muscle tissue was isolated and purified according to a TRIzol extraction method [42]. The RNA concentration was measured using NanoDrop technology. Total RNA samples were treated with DNase I (Invitrogen) to remove any contaminating genomic DNA. The purified RNA was quantified using a 2100 Bioanalyzer system (Agilent, Santa Clara, CA, USA). Isolated mRNA was fragmented with a fragmentation buffer. The resulting short fragments were reverse transcribed and amplified to produce cDNA. Illumina mRNA-seq libraries of the 29 samples were prepared according to the standard high-throughput method. The quality of the cDNA library was assessed by the Agilent Bioanalyzer 2100 system. The library was sequenced with a paired-end (2 × 150 bp) setting using the Illumina X Ten Sequencing System (Illumina, San Diego, CA, USA). The transcriptome data of muscle tissue in 131 hybrid individuals was obtained using DNA nanoball (DNBSEQ-T7) technology according to the standard method [43]. Then, low-quality bases and adapters were trimmed out using Fastp (v. 0.21.0). The high-quality reads were used in the next analyses.

### Detection of gene expression based on mRNA-seq

All mRNA-seq reads of 2nRR, 2nCC, and the hybrids were mapped to the combined nuclear and mitochondrial genomes of *C. auratus* red var. [8] and *C. carpio* [32] using HISAT2 [44] (v. 2.1.0) with default parameters. Then, the mapped files were handled with SAMtools (v. 1.10) [45], while the unique mapped reads were obtained using htseq-count (v. 0.12.4) [46]. The expression value was normalized based on the ratio of the number of mapped reads for each gene to the total number of mapped reads for the entire genome. The transcripts per million (TPM) values were calculated based on the normalized data. These reads in the F_1_ hybrid and four triploids were used to calculate the expression values of the genes in subgenomes R and C [47]. The genes with mapped reads in each sample < 10 and TPM values < 1 were not used in our next analyses. Differential expression analysis was performed using Deseq2 of the R package with the below thresholds: fold changes > 3, *p*-value < 0.001, and Padj < 0.001. Differential expression analysis was performed in 13 individuals of the diploid group (2nRC and 2nCR) and in 10 individuals of the allotriploid group (3nRC_2_ and 3nC_2_R).

Both WGS and mRNA-seq were performed in the 29 individuals (eight months after hatching, 7 in 2nRC, 4 in 3nR_2_C, 7 in 3nRC_2_, 8 in 3nCR_2_, and 3 in 3nC_2_R) for investigating the effects of CNVs on allele-specific expression (ASE) and species-specific expression, which were assessed based on the above thresholds of differential expression analysis.

### Weighted gene correlation network analysis and functional annotation

To investigate expression patterns across samples, we conducted coexpression analysis based on the two F_1_ hybrid and triploid samples using weighted gene correlation network analysis (v. 1.67). An unsupervised network on gene expression was built using the following default parameters: First, a matrix of Pearson correlations between genes was generated based on expression values. Then, an adjacency matrix representing the connection strength among genes was established by raising the correlation matrix to a soft threshold power. Next, the adjacency matrix was used to calculate a topological overlap matrix. Genes with similar coexpression patterns were clustered using hierarchical clustering of dissimilarity. Pearson correlations between the expression level of that gene and module were performed using eigengene-based connectivity, while Pearson correlations were further calculated to measure the strength and direction of association between modules and growth traits. The coexpressed modules were determined and used in our next functional analyses. The hub genes related to growth regulation were further filtered based on thresholds of module membership > 0.8 and gene significance > 0.3. Functional enrichment analyses were conducted and annotated with GO and KEGG databases. The standard deviation of body weight divided by the average body weight across individuals was used to assess the body weight diversity in each population. The standard deviation of the TPM value across the individuals of the six hybrid varieties was used to assess the expression diversity of the predicted growth-regulated genes.

### Correlation in the expression of alleles R and C, correlation between growth traits and allelic expression

We detected a Pearson Correlation Coefficient (PCC) between the expression of alleles R and C based on the below thresholds: a positive correlation was settled as PCC > 0.3; a strongly positive correlation was settled as PCC > 0.5; a negative correlation was settled as PCC < -0.3; a strongly negative correlation was settled as PCC < -0.5. Differential analysis was performed based on the *p*-value using a *t*-test. Then, we performed Pearson’s correlation analysis between growth weight and the expression of allele R or C.

### Mapping of methyl-seq data and differentially methylated analysis

The whole-genome bisulfite sequencing data of 2nRR, 2nCC, 3nR_2_C, and 3nRC_2_ (muscle tissue, 2 years old, three biological replicates in each variety) were obtained from NGDC database (BioProject accession no. PRJCA003625). After quality checking of the methyl-seq reads, the clean reads of 2nRR and 2nCC were mapped to the respective genomes, and the clean reads of the two triploids (3nR_2_C and 3nRC_2_) were mapped to the combined genome sequences of 2nRR and 2nCC [8, 32]. The Bismark analysis pipeline was used to detect the methylated loci with the mapped parameters (-score_min L, 0, −0.2 −X 1000 -no-mixed -no-discordant) [48, 49]. The clean reads were used for mapping to the reference genome four times, and only the reads that mapped to the same position of the reference genome each time were retained in our next analysis. A binomial distribution test was performed to identify 5-methylcytosine for each cytosine site. The potential methylation sites were then checked using the depth > 4 and false discovery rate (FDR) 0.05 thresholds.

The average CpG methylation was detected in different gene regions, including upstream (a window size of 100 bp for 2 kb regions), the gene body, and downstream (a window size of 100 bp for 2 kb regions) of the coding regions. The average CpG methylation in the upstream and downstream transposon regions (2 kb) was calculated and plotted using R. The regions with different methylation were detected using MOABS [50]. The R packages DSS and bsseq were used to call differentially methylated regions and predict differentially methylated genes based on a *p*-value < 0.01.

### Hematoxylin and eosin staining

A 10 mm trunk muscle from 2nRR, 2nCC, 3nR_2_C, and 3nRC_2_ was dissected from the region in the dorsum and fixed in Bouin’s solution for 24 hours. The fixed tissues were washed with distilled water for 4 hours at 20 ℃. After dehydration in ethanol gradients and xylene, the samples were fixed in 4% paraformaldehyde and cut into serial paraffin sections (5–7 μm in thickness). Sections were processed for hematoxylin and eosin (HE) staining. Digital images were captured with a microscope (DX8; Olympus, Tokyo, Japan). Three independent biological replicates were used to collect quantitative data on HE staining in each hybrid variety.

### Animal ethics declarations

All procedures performed on animals were approved by the academic committee at Hunan Normal University, Hunan, China. (Approval number: 2018D013).

## Data availability

The raw reads of WGS and mRNA-seq data have been deposited in the National Genomics Data Center (NGDC) under the BioProject (https://ngdc.cncb.ac.cn/bioproject/browse/PRJCA013677) with accession numbers CRA009160 (WGS data for 29 individuals) and CRA009161 (mRNA-seq data for 29 individuals), and CRA009164 (mRNA-seq data for 131 individuals). The raw reads of equally mixed DNA from 2nRR and 2nCC have been deposited in NGDC (accession number: SAMC449140). The raw reads of DNA methylation have been deposited in NGDC under the BioProject accession number PRJCA003625.

## CRediT author statement

**Li Ren:** Conceptualization, Data curation, Formal analysis, Writing – original draft, Project administration, Writing – review & editing, Validation, Funding acquisition. **Mengxue Luo:** Data curation, Writing – review & editing, Validation. **Jialin Cui:** Extracted the raw material. **Xin Gao:** Data curation, Extracted the raw material. **Hong Zhang:** Extracted the raw material. **Ping Wu:** Helped with the experiment of H&E staining. **Zehong Wei:** Extracted the raw material. **Yakui Tai:** Extracted the raw material. **Mengdan Li:** Extracted the raw material. **Shaojun Liu:** Conceptualization, Project administration, Validation, Funding acquisition.

## Acknowledgements

This research was supported by National Natural Science Foundation of China (32293252, 32341057, U19A2040, and 32273111), Hunan Provincial Natural Science Foundation (2022JJ10035), Huxiang Young Talent Project of China (2021RC3093), National Key Research and Development Plan Program (2023YFD2401602), Laboratory of Lingnan Modern Agriculture Project (NT2021008), Special Funds for Construction of Innovative Provinces in Hunan Province (2021NK1010), Earmarked fund for China Agriculture Research System (CARS-45), and 111 Project (D20007). We extend our gratitude for the valuable assistance provided by English teacher Zhenzhen Dong in professional language editing at Hunan Normal University.

