## Supplementary Figures for "Variation and interaction of distinct subgenomes contribute to growth diversity in intergeneric hybrid fish"

Table of Contents:

Figures S1 to S7	pgs. 2-8

Supplementary Figures

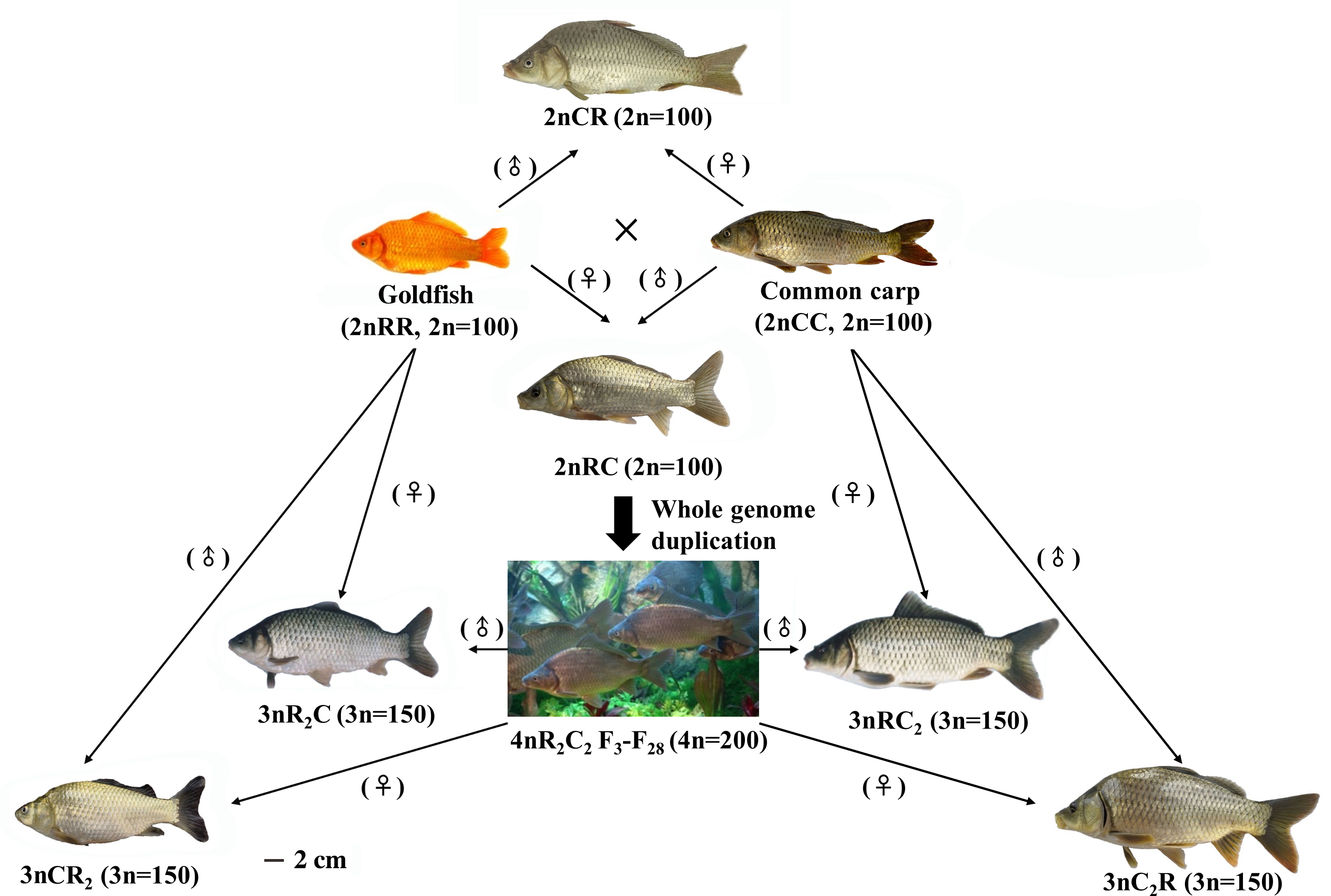

**Figure S1. Appearance of goldfish (2n = 100), common carp (2n = 100), their reciprocal diploid hybrids (2nRC and 2nCR, 2n = 100), their allotetraploid progenies (4nR_2_C_2_, 4n = 200), and the four allotriploid varieties (3nR_2_C, 3nCR_2_, 3nC_2_R, and 3nRC_2_, 3n = 150) derived from interploid crossings with their inbred parents.**

These individuals were bred for 24 months after hatching (average size in each hybrid variety).

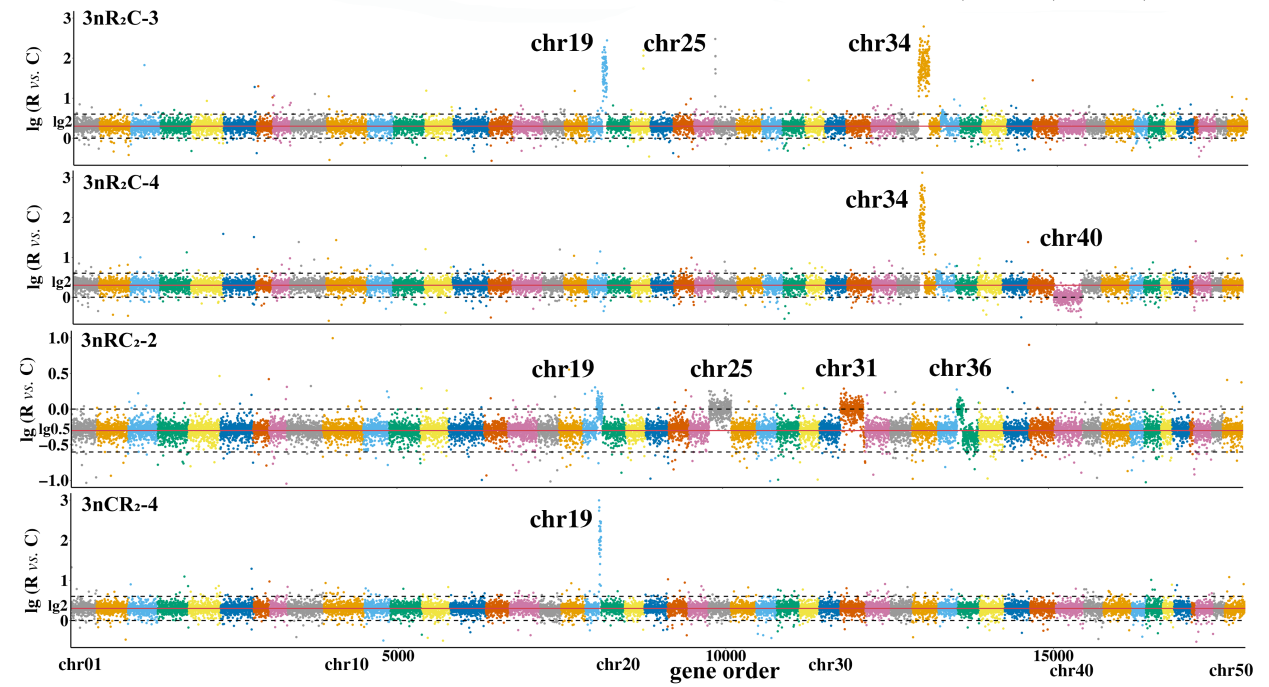

**Figure S2. CNVs in the four triploid individuals.**

The ratio changes of allelic copy number in the four individuals of the triploids. For example, the loss events of whole alleles R or C were observed in the 92 contiguous genes of chr19 and 222 contiguous genes of chr34 in 3nR_2_C-3, the 222 contiguous genes of chr34 in 3nR_2_C-4, and the 92 contiguous genes of chr19 in 3nCR_2_-4. The 1:1 ratio of allelic copy numbers in chr40 of 3nR_2_C-4 and chr19, chr36, chr25, and chr31 of 3nRC_2_-2.

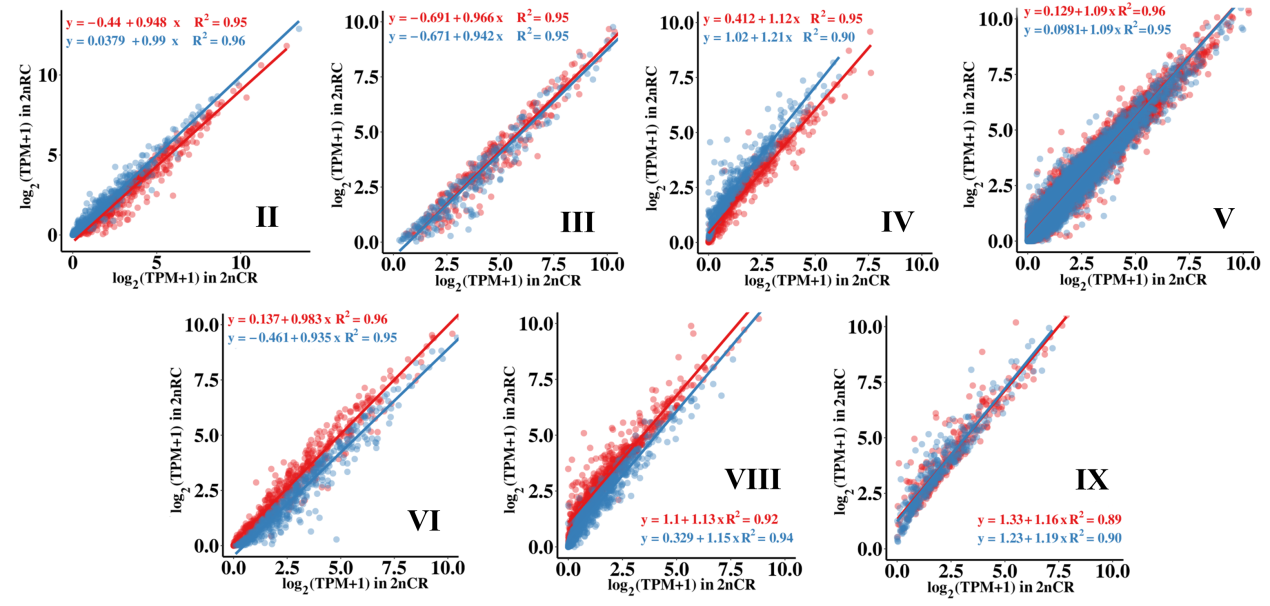

**Figure S3. Distribution of seven allelic expression patterns in reciprocal diploid hybrids (2nRC and 2nCR).** Red represents the expression of allele R, while blue represents the expression of allele C.

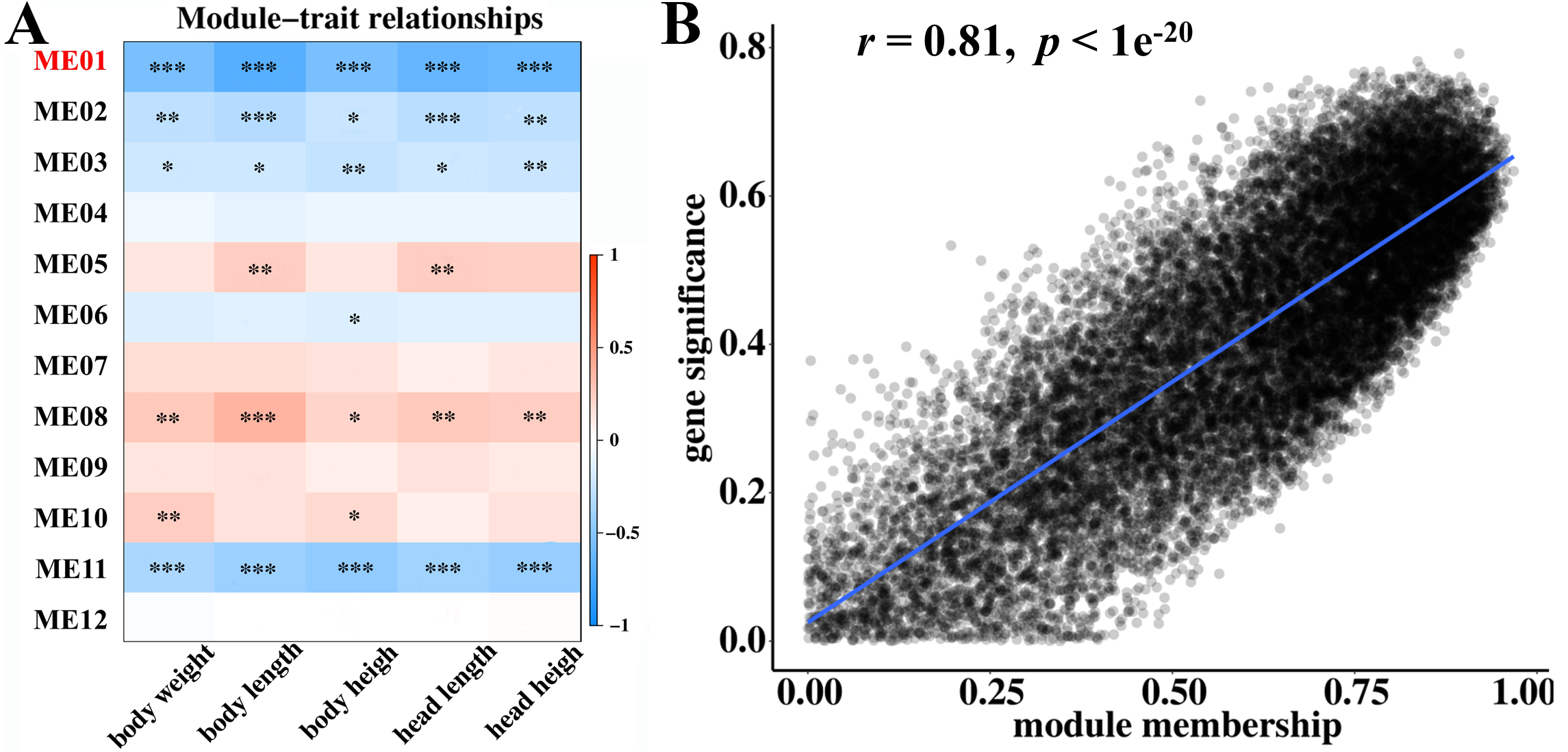

**Figure S4. Gene modules correlated with growth in the hybrid varieties.**

**A.** Correlation between coexpressed modules and sampling traits. Heatmap color represents correlation coefficients (“*” represents Fisher’s asymptotic *p* < 0.05; “**”: *p* < 0.01; “***”: *p* < 0.001). Rows are different modules, and columns are different growth traits. **B.** Distribution of gene significance and module membership of Module Eigengene 1 (ME01).

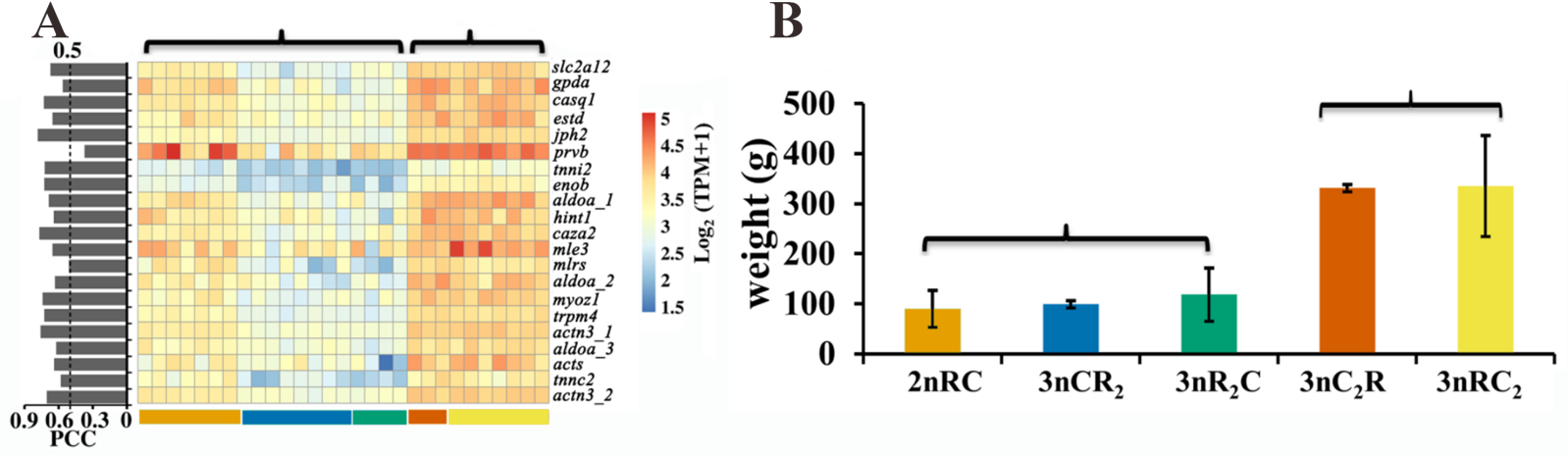

**Figure S5. Correlational analyses between the expression of genes in subgenome C (21 growth-regulated genes) and body weight.**

**A.** Heatmap exhibiting the expression of genes in subgenome C across the 29 individuals (eight months after hatching). The distribution of PCC values in the 21 genes. **B.** The two groups of the five hybrids were classified based on body weight (8 months after hatching) and gene expression.

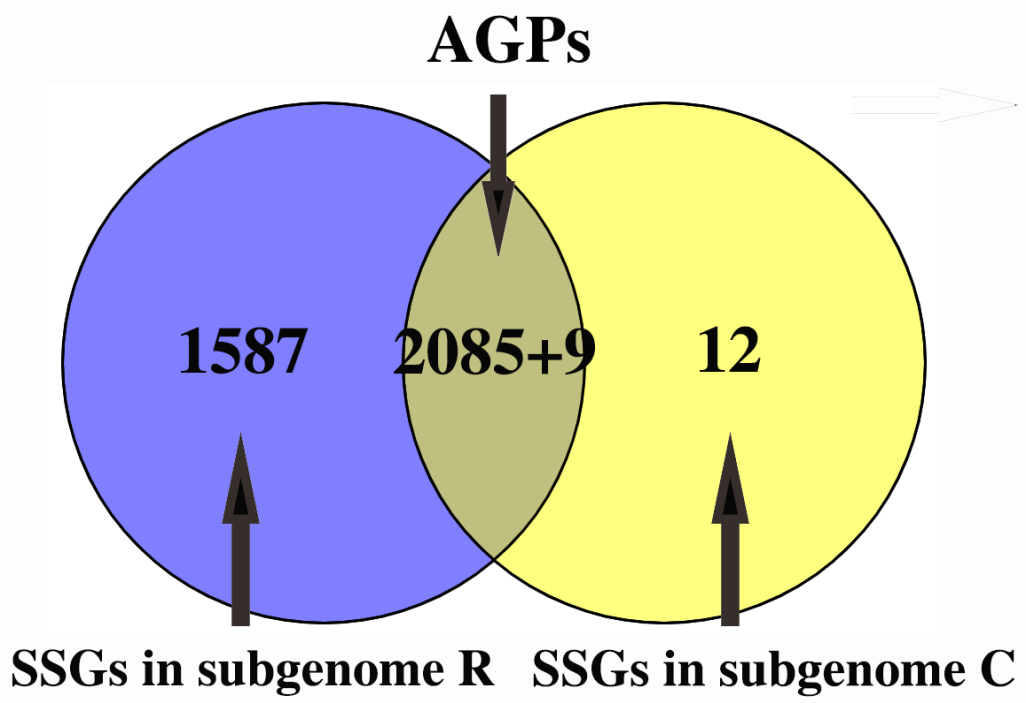

**Figure S6. Distribution of SSGs and AGPs of the 3693 growth-regulated genes.**

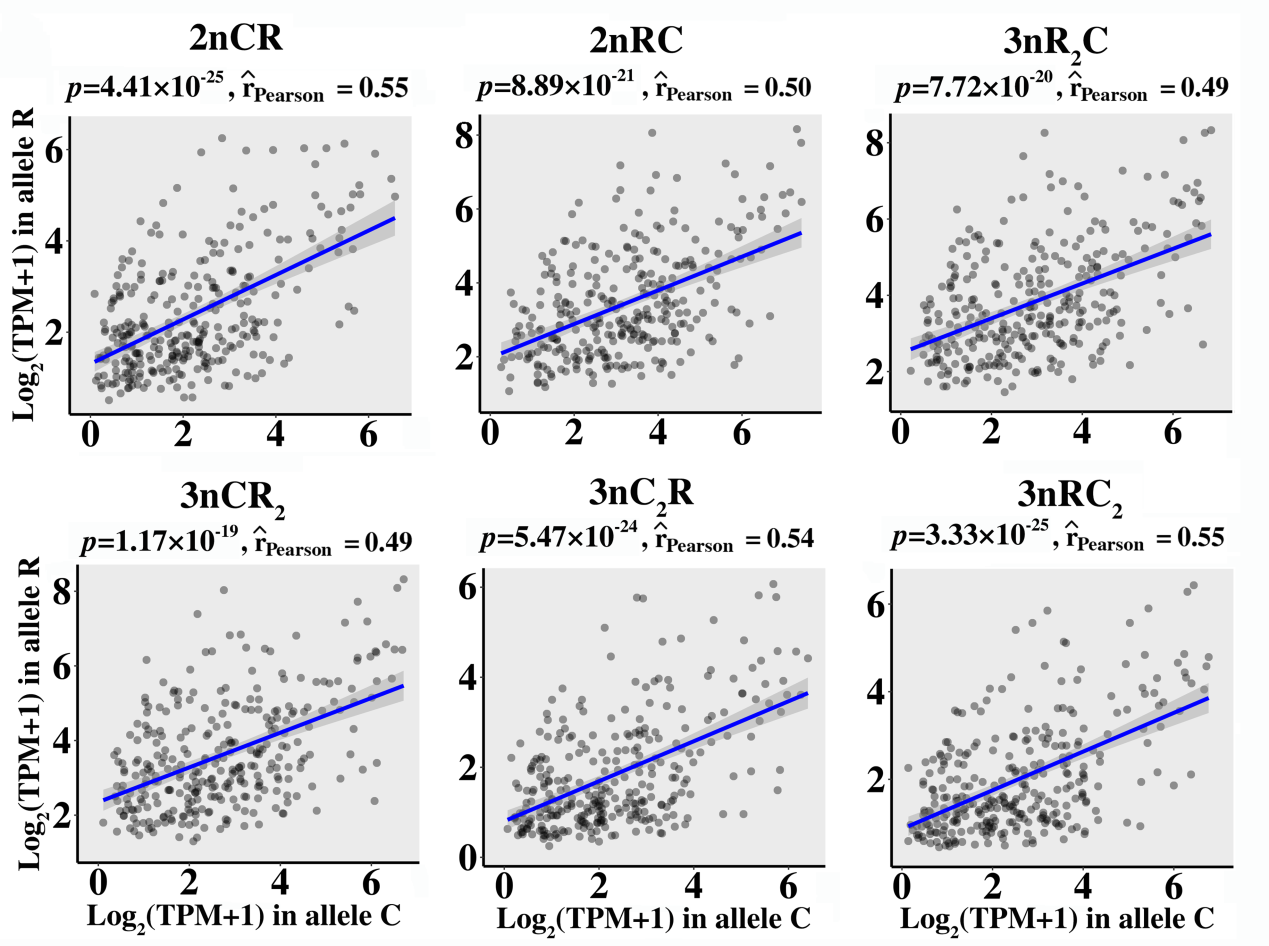

**Figure S7. Growth-regulated genes with strong positive correlation (PCC > 0.5) between expression and body weight.**

833 genes in 2nRC, 812 genes in 2nCR, 1315 genes in 3nR_2_C, 1182 genes in 3nCR_2_, 666 genes in 3nRC_2_, and 1149 genes in 3nC_2_R
