## Supplementary Tables for "Variation and interaction of distinct subgenomes contribute to growth diversity in intergeneric hybrid fish"

Table of Contents:

Tables S1 to S9	pgs. 2-10

**Supplementary Tables**

**Table S1.** Sampling information for the hybrid varieties originating from hybridization of goldfish (2nRR) and common carp (2nCC).

|  |  | mtDNA origin | Subgenome | Ploidy level | No. of individuals and sequencing strategy |
| --- | --- | --- | --- | --- | --- |
| 2nRR | Inbred parent | 2nRR | RR | Diploid |  |
| 2nCC | Inbred parent | 2nCC | CC | Diploid |  |
| 2nRC | Hybrid progeny | 2nRR | RC | Diploid | 7 (WGS) + 7 (8 months, mRNA-seq)+ 33 (24 months, mRNA-seq) |
| 2nCR | Hybrid progeny | 2nCC | RC | Diploid | 13 (24 months, mRNA-seq) |
| 3nR_2_C | Hybrid progeny | 2nRR | RRC | Triploid | 4 (WGS) + 4 (8 months, mRNA-seq)+ 16 (24 months, mRNA-seq) |
| 3nRC_2_ | Hybrid progeny | 2nCC | RCC | Triploid | 7 (WGS) + 7 (8 months, mRNA-seq)+ 43 (24 months, mRNA-seq) |
| 3nCR_2_ | Hybrid progeny | 2nRR | RRC | Triploid | 8 (WGS) + 8 (8 months, mRNA-seq)+ 16 (24 months, mRNA-seq) |
| 3nC_2_R | Hybrid progeny | 2nRR | RCC | Triploid | 3 (WGS) + 3 (8 months, mRNA-seq)+ 10 (24 months, mRNA-seq) |

Note: Whole Genome Sequencing (WGS) was performed on all 29 samples, which were collected 8 months after hatching. This dataset includes 29 samples collected at the 8 months after hatching and 131 hybrid individuals obtained 24 months after hatching.

**Table S2.** Summary of WGS and transcriptome data for 29 individuals.

|  | **WGS** | | | | |  | **Transcriptome** | |
| --- | --- | --- | --- | --- | --- | --- | --- | --- |
| Sample | Total raw data (Gb) | No. of raw reads | Total clean data (Gb) | No. of clean reads | Q20 | Sequencing platform | Total clean data (Gb) | No. of clean reads |
| 2nRCC+2nCC | 133.18 | 887,886,184 | 131.13 | 880,025,966 | 97.07% | Illumina | / | / |
| 3nR_2_C-1 | 16.50 | 109,978,906 | 16.48 | 109,841,182 | 95.48% | Illumina | 7.17 | 23,910,714 |
| 3nR_2_C-2 | 15.90 | 105,983,584 | 15.87 | 105,830,272 | 95.53% | Illumina | 6.53 | 21,754,084 |
| 3nR_2_C-3 | 17.94 | 119,615,940 | 17.92 | 119,494,126 | 95.30% | Illumina | 7.93 | 26,430,006 |
| 3nR_2_C-4 | 20.00 | 133,319,768 | 19.98 | 133,178,110 | 95.41% | Illumina | 6.85 | 22,821,814 |
| 3nRC_2_-1 | 16.99 | 113,251,606 | 16.97 | 113,135,980 | 95.39% | Illumina | 6.88 | 22,943,014 |
| 3nRC_2_-2 | 18.51 | 123,390,314 | 18.49 | 123,248,094 | 95.90% | Illumina | 7.00 | 23,331,026 |
| 3nRC_2_-3 | 25.45 | 169,673,702 | 25.42 | 169,499,982 | 95.77% | Illumina | 6.78 | 22,595,941 |
| 3nRC_2_-4 | 17.23 | 114,867,956 | 17.21 | 114,760,194 | 95.64% | Illumina | 7.04 | 23,479,965 |
| 3nRC_2_-5 | 22.56 | 150,416,302 | 22.54 | 150,279,146 | 95.34% | Illumina | 6.98 | 23,266,850 |
| 3nRC_2_-6 | 21.27 | 141,783,602 | 21.22 | 141,491,116 | 96.53% | Illumina | 6.88 | 22,923,228 |
| 3nRC_2_-7 | 17.00 | 113,328,292 | 16.98 | 113,220,270 | 95.20% | Illumina | 6.89 | 22,982,121 |
| 2nRC-1 | 15.47 | 103,105,580 | 15.43 | 102,869,808 | 96.48% | Illumina | 7.05 | 23,483,488 |
| 2nRC-2 | 18.66 | 124,422,298 | 18.64 | 124,283,194 | 95.89% | Illumina | 6.68 | 22,250,310 |
| 2nRC-3 | 19.33 | 128,870,236 | 19.28 | 128,552,694 | 96.69% | Illumina | 6.76 | 22,544,535 |
| 2nRC-4 | 19.18 | 127,844,980 | 19.14 | 127,608,750 | 96.47% | Illumina | 6.98 | 23,267,993 |
| 2nRC-5 | 16.96 | 113,083,420 | 16.93 | 112,850,426 | 96.75% | Illumina | 7.74 | 25,790,752 |
| 2nRC-6 | 19.90 | 132,678,094 | 19.88 | 132,540,870 | 95.91% | Illumina | 6.86 | 22,858,862 |
| 2nRC-7 | 20.74 | 138,270,420 | 20.72 | 138,122,580 | 96.60% | Illumina | 6.91 | 23,044,206 |
| 3nCR_2_-1 | 16.08 | 107,207,680 | 16.07 | 107,109,188 | 96.00% | Illumina | 6.65 | 22,180,305 |
| 3nCR_2_-2 | 19.66 | 131,088,560 | 19.63 | 130,842,340 | 96.41% | Illumina | 6.94 | 23,119,498 |
| 3nCR_2_-3 | 18.82 | 125,447,802 | 18.8 | 125,305,184 | 94.78% | Illumina | 6.98 | 23,253,547 |
| 3nCR_2_-4 | 18.01 | 120,045,368 | 17.98 | 119,884,758 | 96.09% | Illumina | 6.88 | 22,925,780 |
| 3nCR_2_-5 | 18.98 | 126,504,438 | 18.96 | 126,382,834 | 95.41% | Illumina | 7.52 | 25,069,355 |
| 3nCR_2_-6 | 15.57 | 103,800,324 | 15.55 | 103,685,582 | 95.93% | Illumina | 6.65 | 22,182,434 |
| 3nCR_2_-7 | 16.17 | 107,785,634 | 16.15 | 107,662,336 | 95.91% | Illumina | 7.11 | 23,706,727 |
| 3nCR_2_-8 | 17.98 | 119,862,946 | 17.96 | 119,723,742 | 94.95% | Illumina | 6.70 | 22,340,204 |
| 3nC_2_R-1 | 17.69 | 117,962,178 | 17.67 | 117,820,468 | 95.74% | Illumina | 6.89 | 22,974,313 |
| 3nC_2_R-2 | 19.33 | 128,894,370 | 19.31 | 128,763,682 | 95.61% | Illumina | 6.74 | 22,479,164 |
| 3nC_2_R-3 | 20.51 | 136,710,642 | 20.46 | 136,410,036 | 96.73% | Illumina | 7.56 | 25,192,814 |
| Total | 671.57 | 4,477,081,126 | 668.77 | 4,464,422,910 |  |  | 202.50 | 675,103,050 |

Note: Sample “2nRCC+2nCC” represents the mixed DNA of goldfish and common carp based on equal amounts.

**Table S3.** Genotypes of hybrid fish inferred from whole-genome sequencing data.

|  | Subgenome R | Subgenome C | Actual rate of R *vs.* C^a^ | Predicted rate of R *vs.* C^a^ | mtDNA type |
| --- | --- | --- | --- | --- | --- |
| 2nRC-1 | 5,845,791 | 3,997,525 | 1.46 | 1.00 | 2nRR |
| 2nRC-2 | 7,207,615 | 4,918,920 | 1.47 | 1.00 | 2nRR |
| 2nRC-3 | 7,862,329 | 5,366,176 | 1.47 | 1.00 | 2nRR |
| 2nRC-4 | 7,074,265 | 4,816,736 | 1.47 | 1.00 | 2nRR |
| 2nRC-5 | 7,341,184 | 5,005,697 | 1.47 | 1.00 | 2nRR |
| 2nRC-6 | 6,341,555 | 4,384,756 | 1.45 | 1.00 | 2nRR |
| 2nRC-7 | 7,468,732 | 5,170,741 | 1.44 | 1.00 | 2nRR |
| 3nC_2_R-1 | 4,503,364 | 6,143,998 | 0.73 | 0.50 | 2nRR |
| 3nC_2_R-2 | 4,912,249 | 6,696,865 | 0.73 | 0.50 | 2nRR |
| 3nC_2_R-3 | 5,196,835 | 7,085,952 | 0.73 | 0.50 | 2nRR |
| 3nCR_2_-1 | 9,558,396 | 3,321,818 | 2.88 | 2.00 | 2nRR |
| 3nCR_2_-2 | 7,861,414 | 2,729,295 | 2.88 | 2.00 | 2nRR |
| 3nCR_2_-3 | 8,072,549 | 2,824,330 | 2.86 | 2.00 | 2nRR |
| 3nCR_2_-4 | 8,093,148 | 2,807,790 | 2.88 | 2.00 | 2nRR |
| 3nCR_2_-5 | 9,803,335 | 3,430,761 | 2.86 | 2.00 | 2nRR |
| 3nCR_2_-6 | 9,417,607 | 3,287,606 | 2.86 | 2.00 | 2nRR |
| 3nCR_2_-7 | 8,958,305 | 3,151,472 | 2.84 | 2.00 | 2nRR |
| 3nCR_2_-8 | 9,076,989 | 3,152,754 | 2.88 | 2.00 | 2nRR |
| 3nR_2_C-1 | 8,258,697 | 2,886,939 | 2.86 | 2.00 | 2nRR |
| 3nR_2_C-2 | 7,986,943 | 2,771,951 | 2.88 | 2.00 | 2nRR |
| 3nR_2_C-3 | 9,088,787 | 3,098,996 | 2.93 | 2.00 | 2nRR |
| 3nR_2_C-4 | 10,046,266 | 3,464,362 | 2.90 | 2.00 | 2nRR |
| 3nRC_2_-1 | 4,306,911 | 5,877,241 | 0.73 | 0.50 | 2nCC |
| 3nRC_2_-2 | 6,557,404 | 8,767,749 | 0.75 | 0.50 | 2nCC |
| 3nRC_2_-3 | 4,704,038 | 6,412,842 | 0.73 | 0.50 | 2nCC |
| 3nRC_2_-4 | 4,382,332 | 5,974,122 | 0.73 | 0.50 | 2nCC |
| 3nRC_2_-5 | 5,739,352 | 7,839,805 | 0.73 | 0.50 | 2nCC |
| 3nRC_2_-6 | 5,392,320 | 7,348,412 | 0.73 | 0.50 | 2nCC |
| 3nRC_2_-7 | 4,322,683 | 5,896,524 | 0.73 | 0.50 | 2nCC |

^a^ represents the rate of the average number of the mapped reads in subgenomes R and C.

**Table S4.** Summary of mitochondrial reads mapped to goldfish and common carp genomes in transcriptome data (29 individuals).

|  | Determined mtDNA type | Reference mitochondrial genome | No. of mapped reads | Percent of mapped reads (%) | No. of clean reads |
| --- | --- | --- | --- | --- | --- |
| 2nRC-1 | 2nRR | 2nCC | 24 | < 0.01 | 51,434,904 |
|  |  | 2nRR | 178,557 | 0.35 |  |
| 2nRC-2 | 2nRR | 2nCC | 44 | < 0.01 | 69,061,290 |
|  |  | 2nRR | 268,614 | 0.39 |  |
| 2nRC-3 | 2nRR | 2nCC | 35 | < 0.01 | 63,804,375 |
|  |  | 2nRR | 190,809 | 0.3 |  |
| 2nRC-4 | 2nRR | 2nCC | 41 | < 0.01 | 62,141,597 |
|  |  | 2nRR | 179,104 | 0.29 |  |
| 2nRC-5 | 2nRR | 2nCC | 24 | < 0.01 | 64,276,347 |
|  |  | 2nRR | 189,879 | 0.3 |  |
| 2nRC-6 | 2nRR | 2nCC | 97 | < 0.01 | 56,425,213 |
|  |  | 2nRR | 114,138 | 0.2 |  |
| 2nRC-7 | 2nRR | 2nCC | 305 | < 0.01 | 66,270,435 |
|  |  | 2nRR | 306,842 | 0.46 |  |
| 3nR_2_C-1 | 2nRR | 2nCC | 125 | < 0.01 | 54,920,591 |
|  |  | 2nRR | 223,073 | 0.41 |  |
| 3nR_2_C-2 | 2nRR | 2nCC | 89 | < 0.01 | 52,915,136 |
|  |  | 2nRR | 170,260 | 0.32 |  |
| 3nR_2_C-3 | 2nRR | 2nCC | 12 | < 0.01 | 59,747,063 |
|  |  | 2nRR | 109,152 | 0.18 |  |
| 3nR_2_C-4 | 2nRR | 2nCC | 28 | < 0.01 | 66,589,055 |
|  |  | 2nRR | 127,791 | 0.19 |  |
| 3nRC_2_-1 | 2nCC | 2nCC | 186,882 | < 0.01 | 84,749,991 |
|  |  | 2nRR | 710 | 0.22 |  |
| 3nRC_2_-2 | 2nCC | 2nCC | 172,830 | < 0.01 | 56,567,990 |
|  |  | 2nRR | 634 | 0.31 |  |
| 3nRC_2_-3 | 2nCC | 2nCC | 169,312 | < 0.01 | 61,624,047 |
|  |  | 2nRR | 669 | 0.28 |  |
| 3nRC_2_-4 | 2nCC | 2nCC | 147,749 | < 0.01 | 57,380,097 |
|  |  | 2nRR | 546 | 0.26 |  |
| 3nRC_2_-5 | 2nCC | 2nCC | 193,063 | < 0.01 | 75,139,573 |
|  |  | 2nRR | 781 | 0.26 |  |
| 3nRC_2_-6 | 2nCC | 2nCC | 158,324 | < 0.01 | 70,745,558 |
|  |  | 2nRR | 716 | 0.22 |  |
| 3nRC_2_-7 | 2nCC | 2nCC | 99,945 | < 0.01 | 56,610,135 |
|  |  | 2nRR | 410 | 0.18 |  |
| 3nCR_2_-1 | 2nRR | 2nCC | 50 | < 0.01 | 63,191,417 |
|  |  | 2nRR | 126,357 | 0.2 |  |
| 3nCR_2_-2 | 2nRR | 2nCC | 53 | < 0.01 | 51,842,791 |
|  |  | 2nRR | 159,495 | 0.31 |  |
| 3nCR_2_-3 | 2nRR | 2nCC | 44 | < 0.01 | 53,831,168 |
|  |  | 2nRR | 124,270 | 0.23 |  |
| 3nCR_2_-4 | 2nRR | 2nCC | 41 | < 0.01 | 53,554,594 |
|  |  | 2nRR | 152,257 | 0.28 |  |
| 3nCR_2_-5 | 2nRR | 2nCC | 86 | < 0.01 | 65,421,170 |
|  |  | 2nRR | 179,742 | 0.27 |  |
| 3nCR_2_-6 | 2nRR | 2nCC | 140 | < 0.01 | 62,652,592 |
|  |  | 2nRR | 221,608 | 0.35 |  |
| 3nCR_2_-7 | 2nRR | 2nCC | 71 | < 0.01 | 59,942,379 |
|  |  | 2nRR | 151,320 | 0.25 |  |
| 3nCR_2_-8 | 2nRR | 2nCC | 67 | < 0.01 | 59,861,871 |
|  |  | 2nRR | 114,635 | 0.19 |  |
| 3nC_2_R-1 | 2nRR | 2nCC | 42 | < 0.01 | 58,910,234 |
|  |  | 2nRR | 91,608 | 0.16 |  |
| 3nC_2_R-2 | 2nRR | 2nCC | 139 | < 0.01 | 64,381,841 |
|  |  | 2nRR | 146,707 | 0.23 |  |
| 3nC_2_R-3 | 2nRR | 2nCC | 281 | < 0.01 | 68,205,018 |
|  |  | 2nRR | 231,780 | 0.34 |  |

2nRR in “Determined mtDNA type” represents the most of mapped reads in the mitochondrial genome originating from goldfish.

2nCC in “Determined mtDNA type” represents the most of mapped reads in the mitochondrial genome originating from common carp.

**Table S5.** Transcriptome-based mitochondrial type of hybrid fish (29 individuals).

|  | mtDNA type | TPM in subgenome R^a^ | TPM in subgenome C^b^ |
| --- | --- | --- | --- |
| 2nRC-1 | 2nRR | 66666.67 | 0 |
| 2nRC-2 | 2nRR | 66666.27 | 0.4 |
| 2nRC-3 | 2nRR | 66666.67 | 0 |
| 2nRC-4 | 2nRR | 66666.67 | 0 |
| 2nRC-5 | 2nRR | 66666.53 | 0.14 |
| 2nRC-6 | 2nRR | 66666.3 | 0.36 |
| 2nRC-7 | 2nRR | 66666.23 | 0.44 |
| 3nR_2_C-1 | 2nRR | 66664.39 | 2.27 |
| 3nR_2_C-2 | 2nRR | 66666.62 | 0.05 |
| 3nR_2_C-3 | 2nRR | 66663.81 | 2.85 |
| 3nR_2_C-4 | 2nRR | 66660.49 | 6.18 |
| 3nRC_2_-1 | 2nCC | 12.83 | 66653.84 |
| 3nRC_2_-2 | 2nCC | 7.02 | 66659.65 |
| 3nRC_2_-3 | 2nCC | 3.24 | 66663.43 |
| 3nRC_2_-4 | 2nCC | 12.74 | 66653.93 |
| 3nRC_2_-5 | 2nCC | 6.32 | 66660.35 |
| 3nRC_2_-6 | 2nCC | 4.79 | 66661.88 |
| 3nRC_2_-7 | 2nCC | 11.28 | 66655.39 |
| 3nCR_2_-1 | 2nRR | 66666.11 | 0.56 |
| 3nCR_2_-2 | 2nRR | 66666.63 | 0.04 |
| 3nCR_2_-3 | 2nRR | 66666.16 | 0.5 |
| 3nCR_2_-4 | 2nRR | 66666.52 | 0.15 |
| 3nCR_2_-5 | 2nRR | 66662.44 | 4.23 |
| 3nCR_2_-6 | 2nRR | 66666.61 | 0.05 |
| 3nCR_2_-7 | 2nRR | 66666.62 | 0.05 |
| 3nCR_2_-8 | 2nRR | 66666.59 | 0.08 |
| 3nC_2_R-1 | 2nRR | 66666.4 | 0.27 |
| 3nC_2_R-2 | 2nRR | 66643.14 | 23.53 |
| 3nC_2_R-3 | 2nRR | 66618 | 48.67 |

^a^TPM values of hybrid were calculated from reads mapped to the reference mitochondrial genome of 2nRR.

^b^TPM values of hybrid were calculated from reads mapped to the reference mitochondrial genome of 2nCC.

**Table S6.** Overview of CNVs predicted from WGS data.

|  | No. of genes with CNVs in subgenome R | Ratio | No. of genes with CNVs in subgenome C | Ratio | Total |
| --- | --- | --- | --- | --- | --- |
| 2nRC-1 | 1778 | 74.11% | 621 | 25.89% | 2399 |
| 2nRC-2 | 1069 | 66.52% | 538 | 33.48% | 1607 |
| 2nRC-3 | 1031 | 66.73% | 514 | 33.27% | 1545 |
| 2nRC-4 | 1225 | 67.46% | 591 | 32.54% | 1816 |
| 2nRC-5 | 1116 | 68.72% | 508 | 31.28% | 1624 |
| 2nRC-6 | 1798 | 76.80% | 543 | 23.20% | 2341 |
| 2nRC-7 | 1246 | 68.61% | 570 | 31.39% | 1816 |
| 3nCR_2_-1 | 1355 | 64.74% | 738 | 35.26% | 2093 |
| 3nCR_2_-2 | 1363 | 59.34% | 934 | 40.66% | 2297 |
| 3nCR_2_-3 | 1438 | 61.80% | 889 | 38.20% | 2327 |
| 3nCR_2_-4 | 1398 | 54.25% | 1179 | 45.75% | 2577 |
| 3nCR_2_-5 | 1360 | 64.30% | 755 | 35.70% | 2115 |
| 3nCR_2_-6 | 1357 | 63.20% | 790 | 36.80% | 2147 |
| 3nCR_2_-7 | 1376 | 62.35% | 831 | 37.65% | 2207 |
| 3nCR_2_-8 | 1351 | 64.06% | 758 | 35.94% | 2109 |
| 3nR_2_C-1 | 1641 | 65.43% | 867 | 34.57% | 2508 |
| 3nR_2_C-2 | 1644 | 64.95% | 887 | 35.05% | 2531 |
| 3nR_2_C-3 | 1419 | 49.25% | 1462 | 50.75% | 2881 |
| 3nR_2_C-4 | 1597 | 57.55% | 1178 | 42.45% | 2775 |
| 3nC_2_R-1 | 1887 | 80.92% | 445 | 19.08% | 2332 |
| 3nC_2_R-2 | 1828 | 81.50% | 415 | 18.50% | 2243 |
| 3nC_2_R-3 | 1795 | 83.33% | 359 | 16.67% | 2154 |
| 3nRC_2_-1 | 1822 | 79.98% | 456 | 20.02% | 2278 |
| 3nRC_2_-2 | 1639 | 54.33% | 1378 | 45.67% | 3017 |
| 3nRC_2_-3 | 1846 | 80.58% | 445 | 19.42% | 2291 |
| 3nRC_2_-4 | 1835 | 78.18% | 512 | 21.82% | 2347 |
| 3nRC_2_-5 | 1735 | 81.76% | 387 | 18.24% | 2122 |
| 3nRC_2_-6 | 1754 | 81.62% | 395 | 18.38% | 2149 |
| 3nRC_2_-7 | 1820 | 80.07% | 453 | 19.93% | 2273 |

**Table S7.** Summary of allelic copy numbers inferred from WGS data.

| Sample | High in allele R^a^ | High in allele C^a^ | Total | Loss event of allele R^b^ | Loss event of allele C^b^ | Body weight (g) |
| --- | --- | --- | --- | --- | --- | --- |
| 2nRC-1 | 237 (60.15%) | 157 (39.85%) | 394 | 0 | 0 | 91 |
| 2nRC-2 | 211 (59.60%) | 143 (40.40%) | 354 | 0 | 0 | 125 |
| 2nRC-3 | 181 (56.92%) | 137 (43.08%) | 318 | 0 | 0 | 109 |
| 2nRC-4 | 214 (60.62%) | 139 (39.38%) | 353 | 0 | 0 | 112 |
| 2nRC-5 | 199 (59.58%) | 135 (40.42%) | 334 | 0 | 0 | 123 |
| 2nRC-6 | 274 (68.33%) | 127 (31.67%) | 401 | 0 | 0 | 24 |
| 2nRC-7 | 218 (61.41%) | 137 (38.59%) | 355 | 0 | 0 | 43 |
| 3nC_2_R-1 | 233 (62.30%) | 141 (37.70%) | 374 | 8 | 0 | 325 |
| 3nC_2_R-2 | 201 (65.05%) | 108 (34.95%) | 309 | 3 | 0 | 326 |
| 3nC_2_R-3 | 215 (66.36%) | 109 (33.64%) | 324 | 1 | 0 | 341 |
| 3nCR_2_-1 | 229 (60.74%) | 148 (39.26%) | 377 | 0 | 2 | 137 |
| 3nCR_2_-2 | 222 (56.20%) | 173 (43.80%) | 395 | 0 | 2 | 145 |
| 3nCR_2_-3 | 256 (59.67%) | 173 (40.33%) | 429 | 0 | 3 | 27 |
| 3nCR_2_-4 | 238 (52.65%) | 214 (47.35%) | 452 | 0 | 94 | 165 |
| 3nCR_2_-5 | 218 (59.08%) | 151 (40.92%) | 369 | 0 | 1 | 149 |
| 3nCR_2_-6 | 290 (66.97%) | 143 (33.03%) | 433 | 0 | 6 | 151 |
| 3nCR_2_-7 | 246 (62.76%) | 146 (37.24%) | 392 | 0 | 2 | 27 |
| 3nCR_2_-8 | 266 (64.25%) | 148 (35.75%) | 414 | 0 | 1 | 143 |
| 3nR_2_C-1 | 299 (62.03%) | 183 (37.97%) | 482 | 0 | 3 | 92 |
| 3nR_2_C-2 | 305 (63.02%) | 179 (36.98%) | 484 | 0 | 1 | 100 |
| 3nR_2_C-3 | 405 (63.48%) | 233 (36.52%) | 638 | 0 | 118 | 93 |
| 3nR_2_C-4 | 380 (63.33%) | 220 (36.67%) | 600 | 0 | 223 | 110 |
| 3nRC_2_-1 | 232 (62.87%) | 137 (37.13%) | 369 | 0 | 2 | 394 |
| 3nRC_2_-2 | 518 (74.43%) | 178 (25.57%) | 696 | 1 | 1 | 341 |
| 3nRC_2_-3 | 191 (60.06%) | 127 (39.94%) | 318 | 0 | 1 | 426 |
| 3nRC_2_-4 | 226 (64.20%) | 126 (35.80%) | 352 | 2 | 0 | 355 |
| 3nRC_2_-5 | 195 (67.01%) | 96 (32.99%) | 291 | 2 | 0 | 377 |
| 3nRC_2_-6 | 198 (65.13%) | 106 (34.87%) | 304 | 1 | 0 | 97 |
| 3nRC_2_-7 | 237 (65.65%) | 124 (34.35%) | 361 | 2 | 0 | 355 |

^a^ Except for whole allele loss in a gene, the number of genes with a high copy number in allele R or C when compared to the other.

^b^The number represents that gene copies were only from allele R or allele C.

**Table S8.** Overview of transcriptome data from 131 individuals in the six hybrid varieties.

|  | Total raw data (Gb) | No. of raw reads (million) | Total clean data (Gb) | No. of clean reads (million) | Q20 | Sample number |
| --- | --- | --- | --- | --- | --- | --- |
| 2nRC | 420.09 | 2800.58 | 416.13 | 2794.81 | 96.33% | 33 |
| 2nCR | 137.63 | 917.50 | 136.23 | 914.08 | 96.08% | 13 |
| 3nRC_2_ | 453.63 | 3024.19 | 448.65 | 3012.21 | 96.07% | 43 |
| 3nR_2_C | 192.47 | 1283.11 | 190.39 | 1279.37 | 96.30% | 16 |
| 3nCR_2_ | 201.80 | 1345.35 | 200.01 | 1342.95 | 96.54% | 16 |
| 3nC_2_R | 100.13 | 667.53 | 99.22 | 665.10 | 95.94% | 10 |
| Total | 1505.74 | 10038.26 | 1490.63 | 10008.52 | 96.21% | 131 |

**Table S9.** Summary of gene expression changes regulated by mitochondrial genetics.

|  | Group 1 (3nRC_2_ vs. 3nC_2_R) | | | Group 2 (2nRC vs. 2nCR) | | |
| --- | --- | --- | --- | --- | --- | --- |
|  | orthologous genes | SSGs | Total | orthologous genes | SSGs | Total |
| DEGs in subgenome R | 28 (0.16%) | 27 (0.12%) | 55 (0.14%) | 1834 (10.18%) | 1440 (6.50%) | 3274 (8.15%) |
| DEGs in subgenome C | 24 (0.13%) | 47 (0.19%) | 71 (0.17%) | 1508 (8.37%) | 1734 (7.14%) | 3242 (7.66%) |
| Total | 52 (0.29%) | 74 |  | 3342 (18.55%) | 3174 |  |

SSGs: species-specific genes.
